# A scalable human neuron model of Alzheimer’s disease relevant tauopathy reveals mechanisms linking Tau fibrillization to synaptic dysfunction

**DOI:** 10.64898/2026.09.07.749799

**Authors:** Joanna Lipka, Xiwei Shan, Qiao Zhang, Przemyslaw Dutka, Lilian Phu, Matthew C. Johnson, Tim J. Wendorff, Max Adrian, Anne Biton, Dawei Sun, Meena Choi, John G. Moffat, Alexis Rohou, Alessandro Ori, Celine Eidenschenk, Zhixiang Tong, Casper C. Hoogenraad, Claire G. Jeong, Jasvinder K. Atwal

**Affiliations:** Department of Neuroscience, Genentech, Inc., South San Francisco, CA 94080, USA; Department of Functional Genomics, Genentech, Inc., South San Francisco, CA 94080, USA; Department of Structural Biology, Genentech, Inc., South San Francisco, CA 94080, USA; Department of Proteomic and Genomic Technologies, Genentech, Inc., South San Francisco, CA 94080, USA; Department of Pathology, Center for Advanced Light Microscopy & Electron Microscopy, Genentech, Inc., South San Francisco, CA 94080, USA; Department of Biochemical Cellular Pharmacology, Genentech, Inc., South San Francisco, CA 94080, USA

**Keywords:** Tau aggregation, induced pluripotent stem cell derived human neurons, cryo-electron tomography, phosphoproteomics, MARK2, GSK3, PI3K–mTOR, synaptic dysfunction, Alzheimer’s Disease

## Abstract

Tauopathies, including Alzheimer’s disease, are driven by pathological aggregation of hyperphosphorylated Tau, which disrupts synaptic integrity, impairs neuronal communication, and contributes to cognitive decline. To dissect tauopathy pathogenesis and enable therapeutic discovery, reliable and scalable human iPSC-neuron models are essential. Here, we developed two complementary iPSC-derived neuron models: an endogenous Tau seeding model, in which neurons are challenged with pre-formed Tau fragments that form paired helical filament (PHF)-consistent structures, and a Tau-0N3R overexpression seeding model to accelerate pathology. Both models recapitulate hallmark features of tauopathy, including the progressive formation of intracellular, hyperphosphorylated, sarkosyl-insoluble, and conformationally altered Tau aggregates (AT8, MC1 positive), along with synaptic and neuronal dysfunction. Cryogenic electron tomography (cryo-ET) further revealed the morphology of Tau fibrils within cells, as well as the ultrastructure of Tau fibrils trapping synaptic vesicles *in situ*. Using this platform, we performed integrated phosphoproteomics, high-content screening, and functional validation to identify key pathways driving Tau aggregation. MARK2-mediated phosphorylation within Tau’s microtubule-binding domain emerged as an early trigger of aggregation, confirmed by site-specific mutagenesis. In parallel, small molecules targeting the PI3K/mTOR/GSK3 pathway reduced aggregation and restored synaptic function, with GSK3 inhibition lowering phosphorylation at critical aggregation-driving sites on Tau. Together, these findings establish a physiologically relevant, scalable platform for therapeutic screening that connects Tau seed uptake, site-specific phosphorylation, fibril formation, and synaptic disruption, ultimately identifying mechanistically separable intervention points along the aggregation cascade.

**HIGHLIGHTS:**

- Development of scalable iPSC-neuron models enables tauopathy drug discovery and reconstructs progressive Tau seeding, fibrillization and synaptic dysfunction
- Cryo-ET reveals the ultrastructure of Tau fibrils within human neurons and their accumulation at synapses.
- Temporal phosphoproteomics identifies early modulation of MARK-regulated Tau phosphosites.
- PI3K–mTOR and GSK3 regulate distinct stages of the Tau aggregation cascade.
- Site-specific mutagenesis confirms critical Tau residues required for Tau aggregation.

## INTRODUCTION

Neurodegenerative diseases are characterized by the progressive accumulation and spread of misfolded proteins. The hallmarks of Alzheimer’s disease (AD) pathology are extracellular deposition of amyloid-beta (Aβ) plaques accompanied by the formation of intracellular neurofibrillary tangles (NFTs) composed of hyperphosphorylated and aggregated Tau protein. Tau pathology progresses in a stereotypical pattern suggestive of Tau seed propagation across anatomically connected brain regions^1^. The extent of Tau pathology correlates closely with the severity and progression of cognitive decline, as well as with neuronal loss.^2,3,4^ Two recent large-scale Tau-PET studies confirm that Tau pathology predicts cognitive decline.^5,6^ Furthermore, studies of genetic “escapers” of early-onset familial AD (EOFAD) caused by presenilin 1 or 2 (PSEN1/2) mutations report that reduced Tau pathology accumulation or spread, even in the presence of extensive amyloid pathology, correlates with cognitive resilience,^7,8,9^ suggesting Tau deposition may be a key driver of neurodegeneration in AD. Moreover, Tau aggregates are present in several other tauopathies beyond AD, including progressive supranuclear palsy (PSP), frontotemporal dementia (FTD), Pick’s disease, corticobasal degeneration (CBD), and chronic traumatic encephalopathy (CTE). While cryo-EM studies reveal that different tauopathies harbor distinct Tau filament structures^10^, Tau deposition has been proposed to contribute to neurodegeneration across these diseases.

Under normal physiological conditions, Tau’s primary function is to stabilize microtubules. However, in AD, Tau becomes hyperphosphorylated, leading to its detachment from microtubules, adoption of altered structural conformations, with subsequent aggregation into insoluble deposits. These toxic Tau aggregates are believed to disrupt cellular processes, including axonal transport and synaptic function, ultimately leading to neuronal death.^11,12,13^ Tau pathology alters synaptic structure and function in many ways, likely contributing to cognitive decline in tauopathies. Aggregated or hyperphosphorylated Tau impairs axonal transport of essential synaptic components, including vesicles and mitochondria, leading to disrupted synaptic maintenance and plasticity.^14^ Mislocalized Tau accumulates in dendritic spines, where it disrupts excitatory synaptic transmission by altering receptor composition and postsynaptic signaling.^15,16,17^ Additionally, Tau aggregates compromise mitochondrial function and transport at synapses, reducing ATP availability and calcium buffering necessary for neurotransmission.^18^ Tau-induced activation of stress kinases such as Fyn, GSK3β, and p38 MAPK further dysregulates synaptic signaling and receptor trafficking.^19^ Collectively, these mechanisms drive chronic synaptic stress and loss, which correlates more closely with cognitive decline than neuronal death in Alzheimer’s disease and related tauopathies.^20^

Existing preclinical models used to study Tau pathology include transgenic mice or immortalized cell lines expressing non-AD tauopathy-associated mutations (eg. P301L or P301S). However, these models often fail to fully replicate the human AD disease context, including the human neuronal proteome, progressive development of pathology, propagation of seeding, and the paired helical filament (PHF) fold characteristic of AD Tau.^21^ Furthermore, disease-associated mutations may drive Tau aggregation by altering its subcellular localization and/or promoting fibril nucleation, bypassing molecular events critical for early Tau aggregation. Recently, elegant human induced pluripotent stem cell (iPSC) models utilizing endogenous wild-type Tau and AD-derived seeds have successfully enhanced physiological relevance.^22^ However, patient-derived seeds are limited in availability and exhibit variable seeding potency across individuals, restricting their utility for large-scale applications. Recombinant heparin-induced fibrillar Tau frequently used in seeding studies often lacks rigorous quality control, resulting in aggregates with undefined conformations that typically differ from native PHFs.^23^ Therefore, there is a need for a robust, scalable, physiologically relevant human cellular model that faithfully models tauopathy from seeding through synaptic dysfunction to enable mechanistic studies and high-throughput therapeutic discovery.

We developed a highly characterized and scalable human iPSC-derived NGN2 neuron model of tauopathy featuring fibrillar Tau aggregates. By pairing a wild-type Tau-0N3R overexpression (OE) system with recombinant Tau fibril seeds, we generated a model with robust, accelerated pathology that is amenable to automated screening, cryogenic electron tomography (cryo-ET), and phosphoproteomic analysis. Leveraging this platform, we conducted a multi-modal investigation into the drivers of Tau pathology. We demonstrate that seeding iPSC-derived NGN2 neurons with Tau fibrils induces time-dependent Tau aggregation accompanied by a progressive decline in synaptic integrity. In situ cryo-ET directly visualized intracellular Tau fibrils and revealed their accumulation at synapses, leading to network dysregulation. By integrating temporal phosphoproteomic analysis and high-throughput screening, we identified specific kinases and signaling pathways, most notably MARK2, PI3K–mTOR, and GSK3, as regulators of distinct steps in the aggregation cascade. Site-specific mutagenesis distinguished phosphorylation events that control aggregation from canonical pathological Tau epitopes that arise downstream, while pharmacological perturbation revealed separable mechanisms acting through Tau seed uptake and phosphorylation. Together, these findings establish a scalable framework connecting Tau entry, phosphorylation, fibrillization and synaptic pathology in human neurons for therapeutic discovery.

## RESULTS

### Human iPSC neuron models recapitulate tauopathy with robust wild-type Tau aggregate formation

We set out to establish a human neuronal cell model of Tau aggregation displaying disease-relevant inclusions of wild-type endogenous Tau using a seeding approach. Specifically, we cultured wild-type (WT) human iPSC-derived NGN2 (iNGN2) neurons and seeded them at 14 days in vitro (DIV14) with exogenous Tau fibrils, generated *in vitro* using recombinant truncated Tau fragments (amino acid residues 297-391)^24^ to induce aggregation (Fig. 1A). The recombinant Tau fibril preparation used is heparin-free and in our hands produces a heterogeneous mix of filament structures, including paired helical filaments (PHF) that are structurally similar to Tau PHFs found in AD brain (Supplementary Fig. 1A, B). Seeding activity of the recombinant Tau fibrils was confirmed using a Tau aggregate FRET biosensor cell line^25^ (Supplementary Fig. 1C-E).

**Figure 1.**
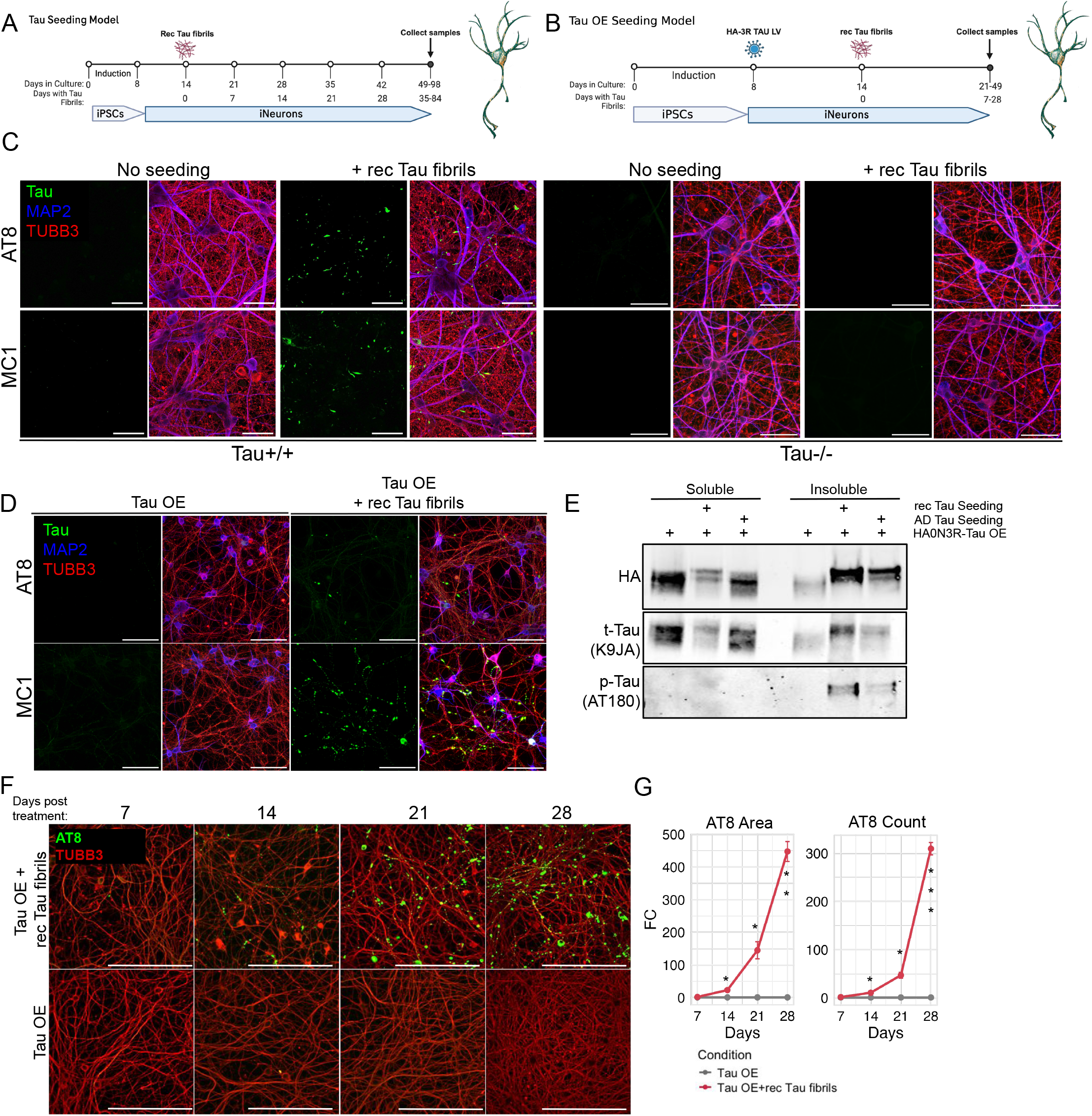
Seeding of iNGN2 Neurons with Recombinant Tau Filaments Induces Aggregation and Hyperphosphorylation of Tau. (**A, B**) Schematic representations of the Tau seeding models. In these models, iPSCs are first differentiated into neurons by induced NGN2 overexpression. In the endogenous seeding model (A), cells are treated with recombinant Tau fibrils one week after replating, and subsequently maintained for 35 to 84 days post-filament treatment to allow aggregate formation. In the Tau OE model (B), one day after replating, neurons are transduced with a lentivirus expressing HA-tagged 0N3R Tau. One week later, the cells are treated with recombinant Tau fibrils and subsequently maintained for 7 to 28 days post-filament treatment to allow aggregate formation. (C) Representative immunofluorescence images of iNGN2 neurons, Tau+/+ (wild type Tau) and Tau-/-(Tau knockout), seeded with recombinant Tau fibrils for 12 weeks as depicted in (A), and stained with AT8 (p-Tau) or MC1 (misfolded Tau) antibodies. Scale bars, 50 µm. (D) Representative immunofluorescence images of iNGN2 neurons overexpressing Tau and seeded with recombinant Tau fibrils for 14 days as in (B), then stained with AT8 or MC1 antibodies. Scale bars, 100 µm. (E) Immunoblot of sarkosyl-soluble and insoluble fractions of lysates from cells overexpressing Tau and seeded with AD Tau or recombinant Tau fibrils, probed with HA, K9JA (total Tau), and AT180 (p-Tau) antibodies. (F) Progressive Tau aggregate formation from 7-28 days. Representative images of iNGN2 neurons overexpressing Tau and seeded with recombinant Tau fibrils for the indicated times and immunostained with AT8 pTau antibody. Scale bars, 200 µm. (G) Quantification of aggregate formation over time. The fold change (FC) in the total area and number of AT8-positive aggregates in seeded versus unseeded cells was calculated. Data represent mean ± SE, n = 3. All comparisons were made relative to the 3R Tau OE (unseeded) condition at the corresponding timepoint. *p<0.05, **p < 0.01, ***p < 0.001 (t-test).

To identify Tau aggregates, we fixed cells at various times after seeding and immunostained them with antibodies recognizing markers of pathological Tau, p-Tau Ser202, Thr205 (AT8)^26^ and MC1 (misfolded Tau).^27^ At DIV49 (5 weeks post-seeding), Tau aggregates were first observed in the axons of iNGN2 neurons seeded with recombinant Tau filaments. Over time, the number of these punctate aggregates gradually increased, and by DIV98 (12 weeks post-seeding), Tau aggregates were clearly detectable (Fig. 1C, Supplementary Fig. 2A). Tau knockout neurons showed no staining with these markers, confirming specific labeling of endogenous, seed-induced aggregates (Fig. 1C). The aggregates were also positive for other p-Tau markers identified in AD brains^28^, including p-Tau Ser409 (PG5), p-Tau Thr231 (AT180), p-Tau Ser422, and p-Tau Thr217 (Supplementary Fig. 2A). These findings demonstrate that human iPSC-derived neurons expressing endogenous Tau, when seeded with recombinant Tau fibrils, develop Tau inclusions that are recognized by multiple markers of AD-associated Tau fibrils. However, these aggregates take a long time to develop, and usually with a relatively limited signal window, making this model unsuitable for biochemical analyses or therapeutic screening.

To promote more widespread and robust aggregate formation in a shorter time, we next developed a wild-type Tau overexpression (OE) seeding model by transducing neurons with a lentivirus expressing HA-tagged 0N3R Tau before seeding with recombinant Tau fibrils (Fig. 1B). WT neurons primarily express the 0N3R Tau isoform, characteristic of early developmental stages and reflecting the immature state of hiPSC-derived neurons.^22^ AD brain-derived Tau fibrils have been shown to efficiently seed 3R Tau expressing cells.^29^ The Tau OE model demonstrated accelerated and more widespread Tau aggregation compared to our endogenous Tau model, with prominent AT8, MC1, MC6, p-Tau Thr217, p-Tau Ser422, and PG5-positive Tau aggregates visible as early as DIV28 (14 days post-seeding) in both the axons and some cell somas (Fig. 1D, Supplementary Fig. 2B). To biochemically confirm the formation of intracellular aggregates, we performed immunoblot analysis on sarkosyl-soluble and -insoluble fractions of cell lysates after seeding. A significant increase in sarkosyl-insoluble Tau species was detected in neurons seeded with recombinant Tau fibrils, as detected by antibodies against total Tau (K9JA), phosphorylated Tau (AT180), and HA-tagged Tau (Fig. 1E). Similar results were seen with neurons seeded with fibrillar Tau isolated from AD brain tissue (AD Tau, Fig. 1E). Time-course analysis of the Tau OE model revealed a progressive increase in both the area and number of AT8-positive aggregates (Fig.1F, G). Thus, complementary endogenous and Tau OE systems capture different experimental windows of the same seeding-dependent process: the endogenous model preserves wild-type Tau at native expression levels, whereas the OE model accelerates fibrillization sufficiently for biochemical, ultrastructural and scalable perturbational studies. We therefore used the Tau OE system for subsequent mechanistic analysis and screening, with key findings validated in the endogenous model.

### Temporal phosphoproteomics identifies early MARK2-associated remodeling of Tau phosphorylation

Hyperphosphorylation of Tau is associated with its aggregation in AD and can affect both its structure and function. In our seeding models, we observed increased phosphorylation of Tau at multiple sites, localized on punctate aggregates. To further characterize early changes in Tau phosphorylation, as well as broader alterations in kinase networks during aggregation, we performed global phosphoproteomic profiling of total cell lysates from recombinant Tau fibril–seeded and unseeded Tau OE human neurons at 3 hours, 7 days, and 21 days post-seeding (Fig. 2A). Using Tandem Mass Tags (TMT)-based proteomics, we quantified 3090 protein groups and 9453 phosphosites across the three time points. Estimated differences of protein abundance were used to adjust phosphoproteomics data and calculate changes of phosphosite modification across conditions (see Methods for details). Seeded neurons displayed marked alterations in the phosphoproteome, visualized as widespread changes in phosphorylation site abundance, with especially pronounced effects on Tau itself (Fig. 2B). Differential abundance analysis revealed altered phosphorylation of kinases previously implicated in Tau regulation (Fig. 2C). Notably, phosphosites corresponding to Microtubule Affinity–Regulating Kinase (MARK) family members, including MARK2 (S545, S376/S388) and MARK1 (S475), displayed increased phosphorylation as early as day 7 post Tau seeding and persisted through later time points. Our analysis identified increased phosphorylation of several canonical MARK2 sites on Tau, including S262, S305, S324, and S356, localized within the microtubule-binding domain (Fig. 2D). Because modification of this region can alter Tau–microtubule interactions, these results identify MARK2-associated phosphorylation as an early molecular event accompanying seed-induced Tau conversion and nominate these residues for functional testing.

**Figure 2.**
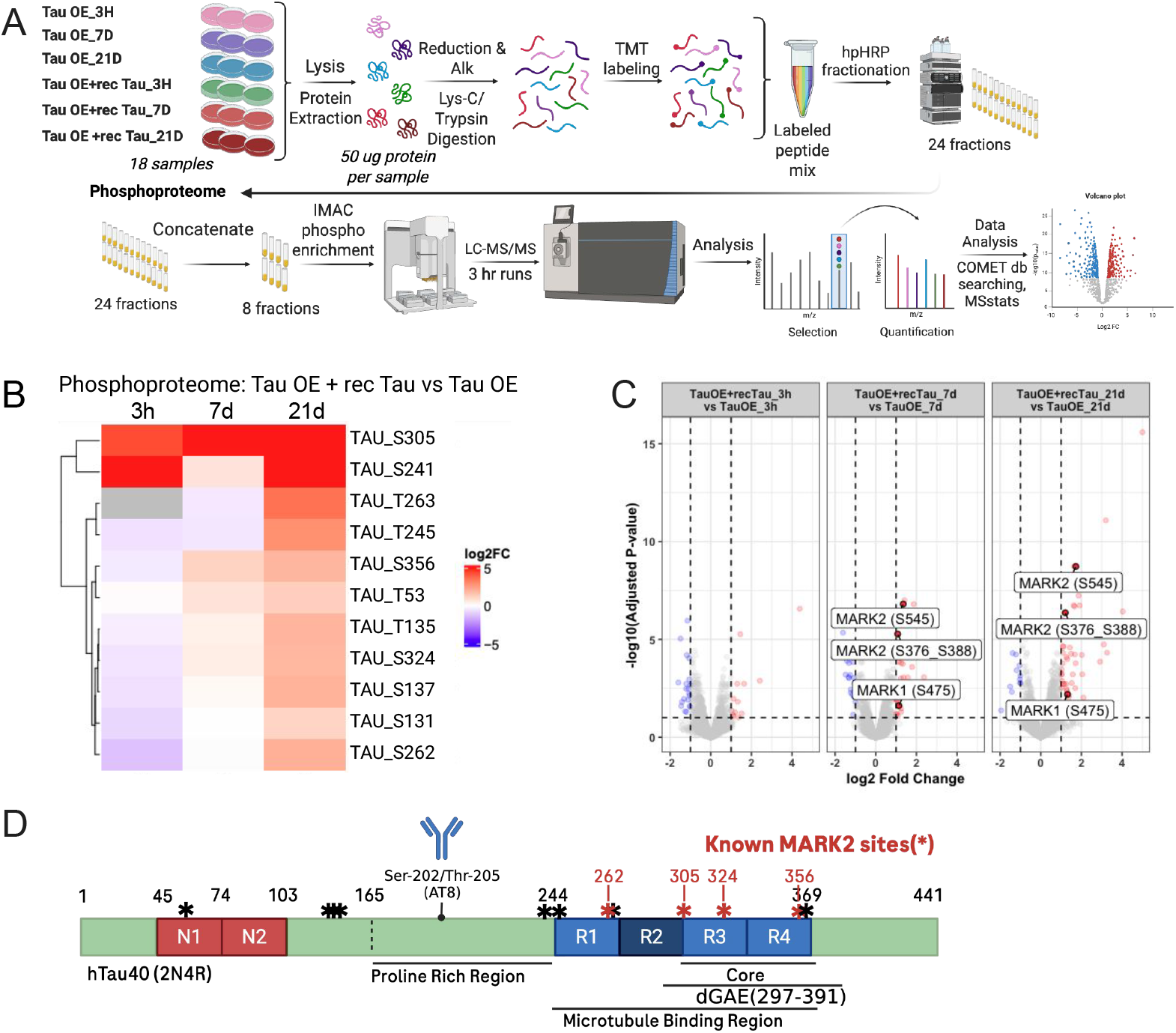
Phosphoproteomic Analysis Reveals Widespread Tau Hyperphosphorylation and Regulation of MARK2 Kinase in Seeded Cells. **(A)** Schematic representation of the mass spectrometry (MS) based phosphoproteomic workflow performed on total cell lysates collected from Tau overexpressing cells seeded with recombinant Tau fibrils for 3 hours, 7 days, or 21 days, and compared to the unseeded control Tau overexpressing cells. **(B)** Heatmap displaying log₂ fold changes in phosphorylation site intensities between seeded and unseeded conditions. Selected Tau sites have an adjusted p-value < 0.05 and a log₂ fold change > 0.58 in at least one of the comparisons (n = 3). **(C)** Volcano plot showing relative changes (log₂ fold) in phosphopeptide abundances, highlighting significant phosphorylation sites in MARK2 and MARK1 kinases. **(D)** Schematic depicting the distribution of Tau phosphorylation sites identified by PTM MS (*), with known MARK2 phosphorylation sites marked in red. Black numbers indicate region boundaries.

### Cryo-ET directly visualizes disease-relevant Tau fibrils within human neurons

To confirm the presence of aggregated Tau in our seeded model and explore the ultrastructural nature of the aggregates, we performed cryo-ET analysis of Tau OE neurons seeded with fibrillar Tau, either recombinant or derived from AD brain tissue. To enable cryo-ET on our cellular tauopathy model, we first developed protocols that allowed us to grow iPSC-derived neurons for an extended period (over six weeks) on electron microscopy (EM) grids (Supplementary Fig. 3A). Tomographic reconstructions of cells 4 hours post-seeding revealed bundles of short extracellular recombinant Tau filaments situated between or above the membranes of neighboring neurites (Fig. 3A; Supplementary Fig. 3C). While these filaments were in close proximity to cellular membranes, no direct contacts were observed. At six weeks post-seeding, we routinely observed multiple examples of intracellular filaments throughout the cytoplasm of cells, though not within membranous compartments. These fibrils exhibited dimorphic morphology, including filaments consistent with the paired helical filaments (PHFs) fold, with a crossover of ∼80 nm, and bundles of flat fibrils, a structure also seen in the seeding material, but much longer (Fig.3B, C; Supplementary Fig. 3D). In contrast, AD Tau seeding yielded a morphologically homogeneous population of intracellular fibrils consistent with PHFs fold (Fig. 3D). These results highlight our model’s ability to capture fold-specific aggregation patterns.

**Figure 3.**
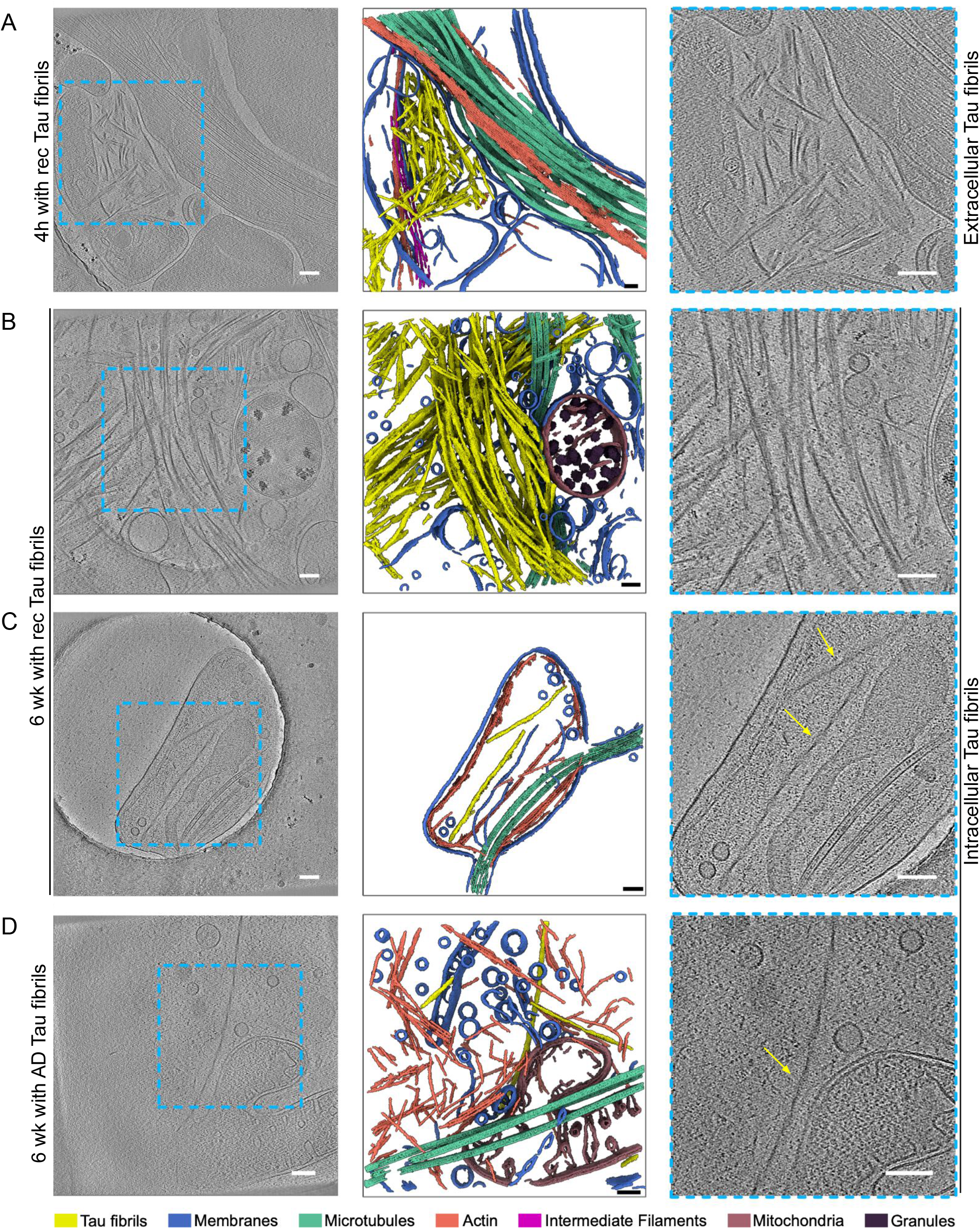
Cryo-ET of Tau Fibril Morphology and Organization in Seeded Cells. Left panels: Central slices from representative tomograms. Middle panels: Segmented volume renderings. Right panels: Enlarged views of the regions highlighted by blue boxes in the left panels, illustrating fibril morphology. (**A**) Extracellular fibrils are observed 4 hours post-seeding with recombinant Tau fibrils. (**B**) Predominant and (**C**) minor (PHF-consistent) intracellular fibril morphologies observed 6 weeks post-seeding with recombinant Tau fibrils. (**D**) Distinct homogeneous (PHF-consistent) fibril morphology observed intracellularly 6 weeks post-seeding with AD Tau. Scale bars, 100 nm.

### Tau aggregation disrupts synaptic structure and function

Given the strong link between Tau pathology and synaptic dysfunction,^30,20^ we next examined whether Tau aggregation compromises synaptic integrity in our seeding models. Immunofluorescence staining of neurons in the Tau OE model seeded with recombinant Tau fibrils revealed pronounced abnormal clustering of presynaptic markers into large foci, including synaptophysin, synapsin, and synuclein, while overall neuronal density (DAPI count) remained largely unchanged (Fig. 4A, B; Supplementary Fig. 4A). Clustering of these synaptic proteins was evident by day 14 post-seeding, coinciding with robust expression of synaptic markers in these cultures. These findings suggest that Tau aggregation induces a selective disruption of synaptic architecture. TUBB3 staining, which labels the neuronal microtubule cytoskeleton and serves as a marker of overall neuronal morphology and health, was unchanged, indicating that neuronal numbers and gross neurite outgrowth were preserved despite Tau-induced synaptic alterations. One caveat is that the abundant 3D growth of TUBB3-positive processes may limit its sensitivity as a readout of axonal integrity. Over time, some signs of neuronal stress became more apparent in the cultures, including reduced ATP levels (CellTiter-Glo), reduced MAP2-positive dendritic area, and accumulation of p62 puncta, suggestive of impaired autophagy and proteostasis.

**Figure 4.**
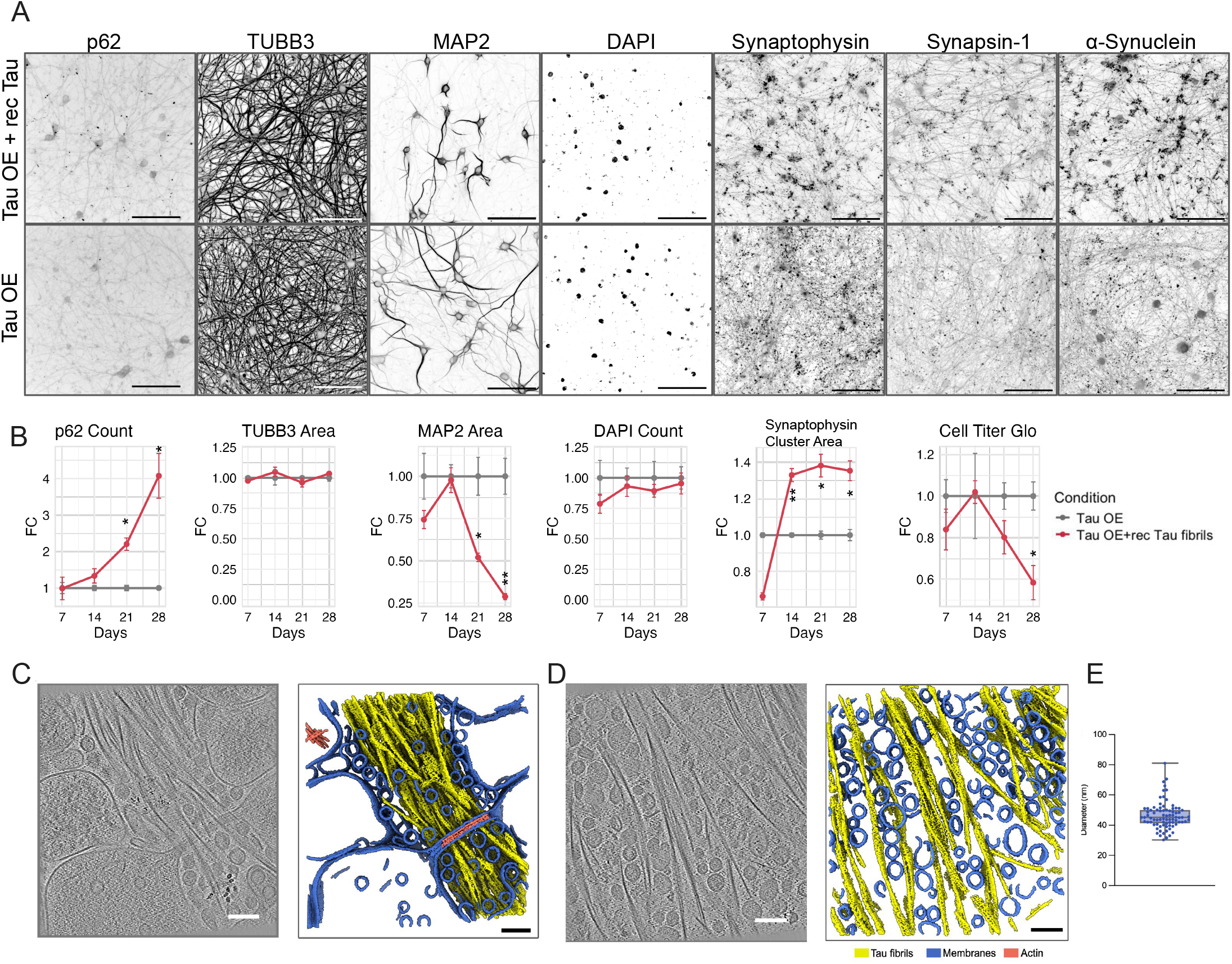
Tau Seeding Induces Progressive Dysfunction of Synaptic Vesicles and Other Markers of Neuronal Integrity. (A) Representative immunofluorescence images of iNGN2 neurons overexpressing Tau and seeded with recombinant Tau fibrils for four weeks, stained for p62, TUBB3, MAP2, DAPI, synaptophysin, synapsin and α-synuclein. Scale bars, 100 µm. (B) Quantification of fold changes (FC) in marker expression comparing seeded versus unseeded cells over time. Data represent mean ± SE, n = 2-5. All comparisons were made relative to the 3R Tau OE condition (unseeded) at the corresponding timepoint. *p<0.05, **p < 0.01, ***p < 0.001 (t-test). (C, D) Visualization of synaptic vesicles associated with Tau fibrils in neurons 6 weeks after seeding with recombinant Tau fibrils. Left panels: Central slices from representative tomograms. Right panels: Corresponding segmented volume renderings. Scale bars, 100 nm. (C) Accumulation of seeding-induced Tau fibrils within synapses. (D) Extracellular seeding-induced Tau fibrils with synaptic vesicles trapped between them. (E) Measured diameters of vesicles trapped between Tau fibrils.

To better define the ultrastructural basis of the observed synaptic disruption, we turned to our cryo-ET analysis. Strikingly, we observed Tau fibrils that accumulate near synaptic vesicles and appear to trap them (Fig. 4C-D). The trapped vesicles could be seen both intracellularly within synaptic regions (Fig. 4C), as well as in what appeared to be extracellular areas, presumably resulting from cytoplasmic membrane rupture, leaving behind "ghost" synapses—similar to how "ghost tangles" are observed in postmortem tissues (Fig. 4D). Quantitative analysis confirmed that the Tau fibril-associated vesicles fell within the expected diameter range of synaptic vesicles (Fig. 4E). Finally, to assess the functional consequences of this pathology at the network level, we performed microelectrode array (MEA) recordings in wild-type seeded versus non-seeded neurons (Supplementary Fig. 4C). Neurons seeded with recombinant Tau fibrils displayed a reduction in the percentage of active electrodes compared to their non-seeded controls, indicating impaired neuronal firing and network connectivity. Similarly to the OE seeding model, in this wild-type model we also observed clustering of synaptic vesicle markers (Supplementary Fig. 4B). Together, these findings demonstrate that Tau aggregation in human iPSC-derived neurons not only reproduces ultrastructural features of disease-relevant Tau filaments, but also disrupts synaptic architecture and impairs neuronal network activity. These results reveal a structural and functional connection between Tau aggregation and synaptic dysfunction in this model, linking synaptic Tau fibril accumulation and vesicle trapping with impaired neuronal network activity. Together, these findings establish a reproducible and scalable platform to investigate disease mechanisms, genetic perturbations, and small-molecule therapeutic interventions.

### High-throughput screening reveals diverse pathways regulating Tau aggregation

We next established a medium throughput drug screening platform to identify small molecules that mitigate Tau aggregation (Fig. 5A, C). We adapted the Tau OE seeding model to 384-well format with automation. We screened a library of approximately 800 well-annotated compounds targeting a variety of different classes of proteins, with a large fraction targeting kinases (Fig. 5B). Cells were treated with compounds at a single dose (3 µM) concurrent with fibril seeding. Based on pilot testing, this dose was found to produce robust effects on Tau aggregation for several compounds without overt toxicity. Compounds were replenished every 4-5 days with media changes, and cells fixed 14 days after fibril treatment to assess Tau aggregation, at a timepoint where there was no overt neurodegeneration (MAP2 Area) and there was already a very good assay window for aggregation (Figure 1 and 4). Hits were defined by their ability to reduce AT8-positive aggregate area (normalized to the live cell number) while preserving cell viability. Compounds were classified as toxic when the MAP2-positive area was less than 60% of the DMSO control (Fig. 5D). As a positive control, we treated cells with Anle138b, a reported inhibitor of Tau, prion, and α-synuclein aggregation.^31,32^ At 10 µM, Anle138b robustly reduced Tau aggregates by ∼50%, validating the assay window (Fig 5E). From the primary screen, we identified approximately 80 compounds that either reduced (“reducers”) or enhanced (“boosters”) Tau aggregation (Fig. 5F, G). Because our primary interest was in compounds that decrease Tau aggregation, the top reducer hits were validated using a nine-point dose–response curve (Fig. 5H, Supplementary Table 1). To confirm that these compounds were specifically modulating Tau aggregation rather than simply reducing Tau phosphorylation (as AT8 recognizes phospho-Tau), we immunostained treated neurons with the conformation-specific antibody MC1, which selectively detects pathological aggregated Tau. Notably, 80% of the hits were confirmed (Supplementary Table 1). Pathway-level analysis of the validated hits revealed a broad distribution across multiple signaling networks (Fig. 5I). Key nodes included the PI3K–AKT axis (PI3K, AKT, mTOR, GSK3, PIM), cell-cycle regulators (Aurora A/B, TTK, Chk1, CDKs), MAPK signaling (ERK1/2, JNK1/3, ASK1, DUSP1/6, PAK1/2/4/5), and NF-κB signaling (IKKβ, NFKB, STAT3). Additional clusters of hits mapped to receptor tyrosine kinases, TGF-β signaling, apoptosis regulators, epigenetic modifiers, immune pathways, and the unfolded protein response (UPR). This diverse spectrum of pathways highlights the complexity of Tau aggregation as a cellular process, suggesting that multiple signaling cascades converge to regulate Tau misfolding, aggregation, and toxicity.

**Figure 5.**
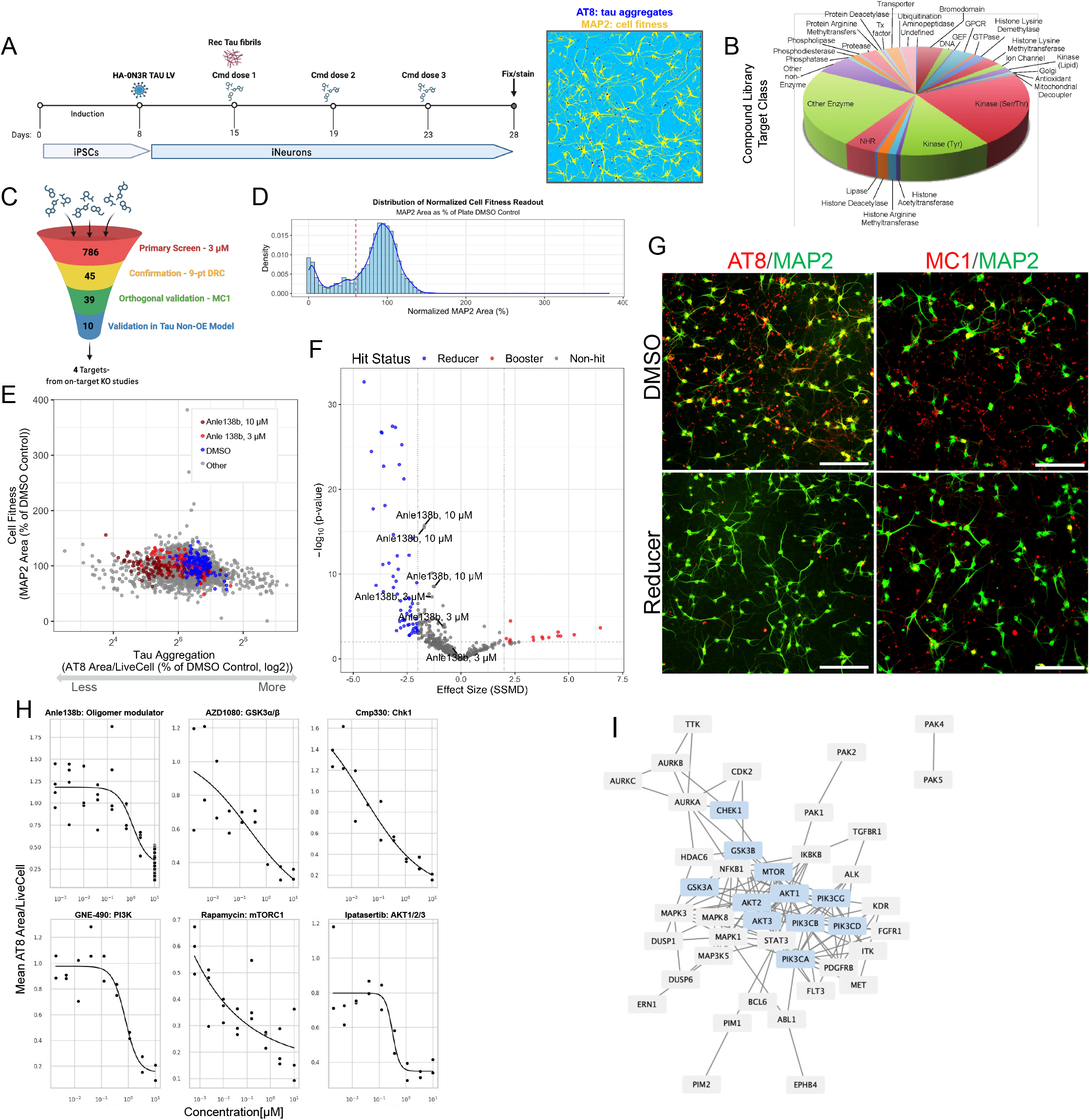
Compound Screen for Small Molecule Reducers of Tau Aggregates. (A) Schematic overview of the primary high-content screening assay. iNGN2 neurons were plated in a 384-well format, transduced with a 0N3R Tau virus, and seeded with recombinant Tau fibrils one week later. The first compound dose was added at the time of fibril treatment, with subsequent doses replenished every four days. Cells were fixed on Day 28 (14 Days after recombinant Tau fibril addition). Samples were stained for AT8 (to assess Tau aggregation efficacy), MAP2 and DAPI (as cell fitness readouts), and then imaged. (B) Classification of the compound library by target class. (C) Schematic representation of the screening funnel from primary hit identification to follow-up validation. The number of compounds (reducers) selected at each step is shown. (D) Distribution of normalized cell fitness readout. Compounds were classified as toxic when the MAP2-positive area was less than 60% of the DMSO control. (E) Scatter plot showing Tau aggregation efficacy versus cell fitness, both normalized to the DMSO control for each plate. Compounds with a MAP2-positive area less than 60% of the DMSO control were not plotted; n>=4. Blue dots, DMSO controls; red dots, Anle138b positive control. (F) Volcano plot displaying statistical significance (–log₁₀ p-value) versus strictly standardized mean difference (SSMD; effect size) for each compound that was not considered toxic. Hits were called when p-value < 0.01 with SSMD < –2 for reducers (marked in blue) and SSMD > 2 for boosters (marked in red). All selected reducers of Tau aggregation exhibited an even stronger effect size than our positive control, Anle138b. (G) Representative immunofluorescence images of cells treated with a screen-identified hit compound, stained with AT8, MC1, and MAP2 antibodies. Scale bars, 200µm. (H) Example 9-point dose–response curves for selected compounds; n >=2. (I) STRING network visualization of protein targets whose modulation reduced Tau aggregation in the primary screen. Highlighted in blue are the hit targets from (H).

### PI3K/mTOR/GSK3 signaling is a central regulator of Tau aggregation

To further validate these compounds as modulators of Tau aggregation, we assessed them in our wild-type seeding model where only endogenously expressed Tau is present (Fig. 6A). In this system, inhibitors of PIM kinase, PI3K, mTOR, Chk1, AKT, and NF-κB signaling emerged as the strongest modulators of Tau aggregation (Fig. 6B, C). Importantly, several compounds, including Rapamycin (mTOR inhibitor), GNE-490 and AZD8835 (PI3K inhibitors), Ipatasertib (AKT inhibitor), and AZD1080 (GSK3 inhibitor), not only reduced MC1-positive Tau inclusions after seeding, but also prevented the synaptic clustering defect revealed by synapsin immunostaining (Fig. 6D, E). These findings established a functional connection between reduced Tau pathology and restored synaptic organization. To confirm the target specificity of these compounds in our model system, we used genetic approaches. Knockout of PI3KCA, GSK3B, MTOR, and CHEK1 reduced Tau aggregation after seeding, confirming the involvement of these kinases in Tau aggregation (Fig. 6F, G). Further genetic dissection of the mTOR pathway revealed that loss of upstream repressors TSC1/TSC2 increased Tau aggregation, while knockout of Raptor or Rictor decreased aggregation, implicating both mTORC1 and mTORC2 in regulating pathology (Fig. 6H).

**Figure 6.**
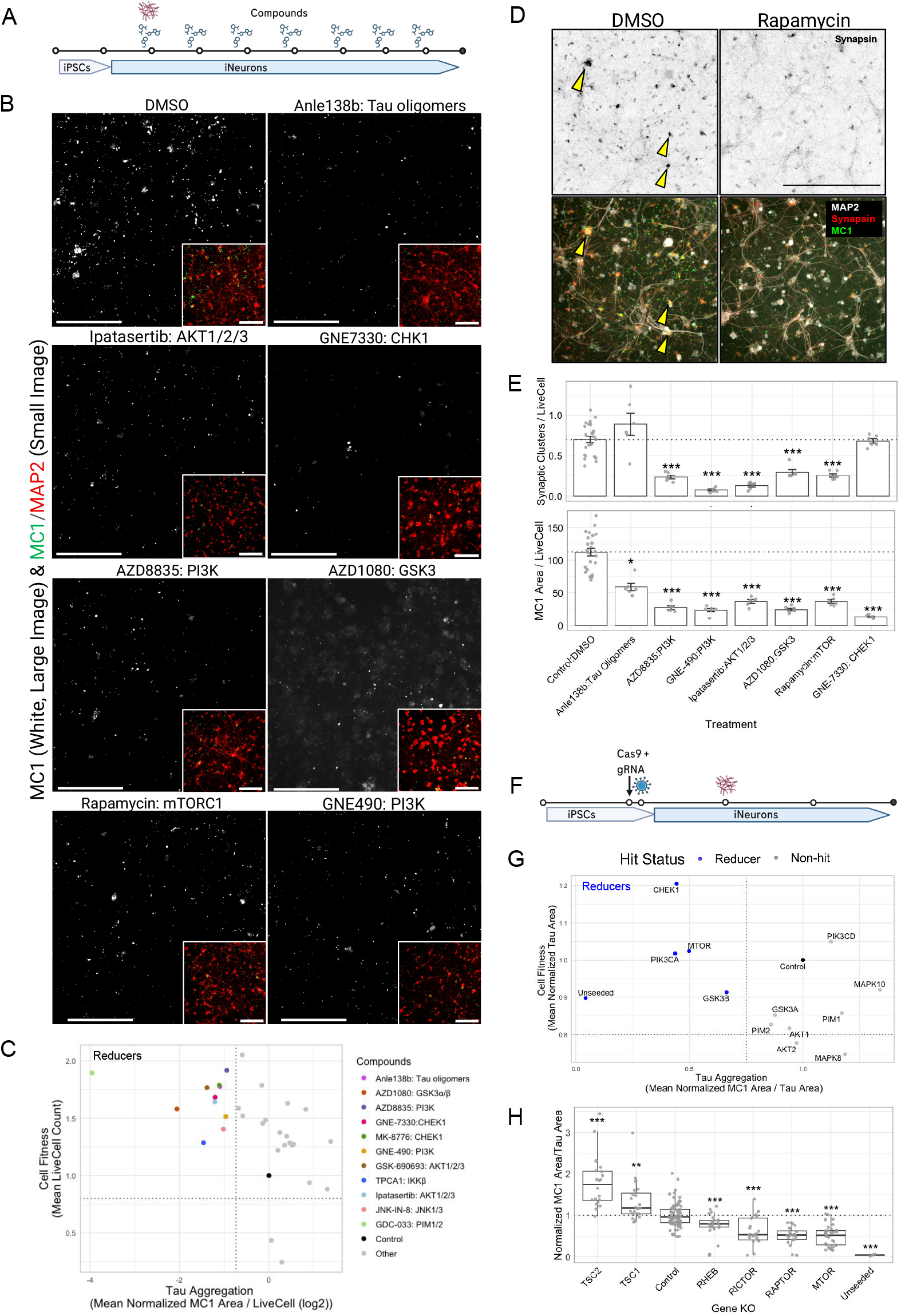
Validation Screens for Small Molecule Reducers of Tau Aggregates. (A) Schematic overview of the validation assays performed in wild-type neurons (B–E). WT iNGN2 neurons were seeded with recombinant Tau fibrils at day 14 and co-treated with compounds. Compounds were applied at the IC50 determined from experiments in Tau OE cells (Table1), and replenished weekly during media changes. Cells were assayed 6 weeks post seeding. (B) Representative immunofluorescence images of iNGN2 neurons treated with select compounds identified in the primary screen, stained for MC1 (white, green in inset) and MAP2 (red in inset). Scale bars, 200 µm. (C) Scatter plot showing Tau aggregation efficacy versus cell fitness for tested compounds, both normalized to the DMSO control for each plate. Hits defined as reducers, were showing at least a 40% reduction in Tau aggregation and no more than a 20% reduction in cell fitness (marked in color). n = 6–12. (D) Representative immunofluorescence images of iNGN2 neurons seeded with recombinant Tau fibrils and co-treated with rapamycin and stained for Synapsin, MC1, and MAP2. Arrows: synaptic clusters. Scale bar, 200 µm. (E) Quantification of synaptic cluster number per live cell (upper panel) and Tau aggregation area per live cell (lower panel) under the treatment of selected compounds. Dotted lines represent DMSO control. Data represent mean ± SEM, n= 5-24. *p < 0.05, ***p < 0.001 (F) Schematic overview of the assay used to confirm the specificity of the selected compounds by genetic knockdown (G, H). iNGN2 neurons were electroporated with RNPs containing Cas9 and the targeting guide RNA on day 8 (at replating), and transduced with HA-0N3R Tau LV one day later. Cells were then seeded with recombinant Tau fibrils one week after viral transduction and fixed on day 28 (14 days after seeding with fibrils). (G) Scatter plot of Tau aggregation efficacy versus cell fitness for each of the genes targeted, both normalized to the non-targeting guide control for each plate. Hits defined as reducers, were showing at least a 25% reduction in Tau aggregation and no more than a 20% reduction in cell fitness (marked in blue). n = 12-80. (H) Quantification of Tau aggregation efficacy, normalized to the non-targeting guide control (OR51S1) for each plate. Genes in the mTOR pathway were probed. Boxplots show the distribution (median, quartiles, spread) of normalized MC1 area values for each gene KO group; n = 20-80. The dotted line indicates control levels. All comparisons were made relative to the control condition following one-way ANOVA with Dunnett’s post hoc test. **p<0.01, ***p<0.001.

These compounds may reduce aggregated Tau levels in cells by altering various different cellular mechanisms, including fibril uptake/seeding, Tau expression, Tau phosphorylation, fibril assembly, or fibril degradation (Fig. 7A,^33^). To investigate fibril uptake, we generated a FRET-based Tau aggregation sensor in Tau knockout iNGN2 neurons, which monitors early seeding events (Fig. 7B). PI3K/mTOR inhibitors markedly reduced the FRET signal, indicating decreased internalization of Tau seeds or reduced initiation of aggregation (Fig. 7C, D). We used a HiBit-tagged Tau reporter line to assess whether any of the compounds altered Tau protein expression. Although some inhibitors (Rapamycin, AZD8835, GNE-490, Ipatasertib) caused a modest 20-30% reduction in Tau expression (Supplementary Fig. 5A, B), this was not enough to explain the much larger 60-80% reduction in Tau aggregates observed for these compounds (Fig. 5H). Together, these results demonstrate that the PI3K/mTOR/GSK3 signaling axis is a central early regulator of Tau aggregation and that pathway inhibition not only suppresses Tau pathology but also preserves synaptic structure, highlighting a mechanistic link with therapeutic potential.

**Figure 7.**
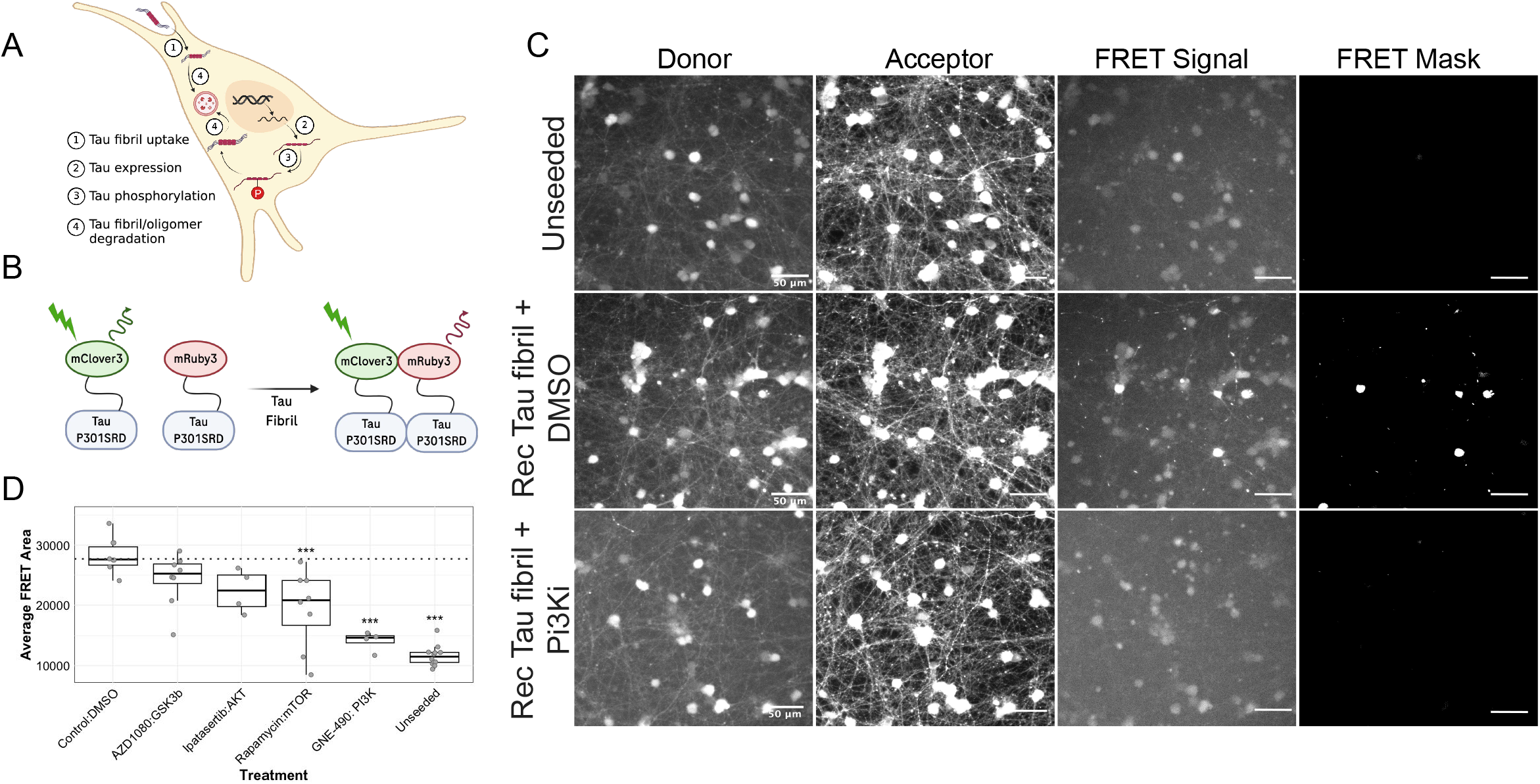
PI3K and mTOR Regulate Uptake of Tau Fibrils in Seeded Cells. (A) Schematic cartoon illustrating the major pathways by which Tau aggregation may be modulated in cells. (B) Schematic representation of the FRET-based Tau aggregation sensor expressed in iNGN2 neurons. Tau-knockout (KO) iNGN2 neurons were transduced with lentiviral vectors encoding P301S^RD–mClover3 (donor) and P301S^RD–mRuby3 (acceptor). Upon addition of Tau fibrils, a FRET signal is detected and quantified. (C) Representative immunofluorescence images of neurons seeded with recombinant Tau fibrils and treated with selected compounds or vehicle control for 3 days. Scale bars, 50 µm. (D) Quantification of the FRET signal. Boxplots show the distribution (median, quartiles, spread) of the average FRET area, n = 4-12. The dotted line indicates control signal from seeded cells treated with DMSO. One-way ANOVA with Dunnett’s post hoc vs control (DMSO). *p < 0.05, **p < 0.01, ***p < 0.001.

### Phosphorylation at GSK3 and MARK2 sites drives Tau aggregation

By integrating phosphoproteomics, high-content screening, and functional validation in the endogenous Tau seeding model, we prioritized two kinase pathways as key drivers of Tau aggregation: the PI3K/mTOR/GSK3 axis and MARK2-mediated phosphorylation. mTOR, GSK3β and MARK2 have been reported to directly phosphorylate Tau^34,35,36,37,38^ and these phosphorylation events have been suggested to promote Tau aggregation. To investigate this connection in our model, we first tested whether pharmacological inhibition of mTOR or GSK3 influences Tau phosphorylation. Acute treatment of seeded Tau OE neurons with AZD1080 (a GSK3 inhibitor) or Rapamycin (an mTOR inhibitor) revealed distinct effects. While mTOR inhibition did not significantly alter Tau phosphorylation at the sites examined, GSK3 inhibition led to a clear reduction in phosphorylation at canonical GSK3 sites on Tau (T217, T231, and S396) (Fig. 8A–C). This suggested that GSK3, but not mTOR, exerts a direct effect on Tau phosphorylation. To investigate a causal relationship between phosphorylation and aggregation, we generated Tau mutants targeting key phosphorylation sites. Sites phosphorylated by GSK3 (T217, T231, S396) and MARK2 (S262, T356-—the most widely reported MARK2 sites^35,39^ identified in our phosphoproteomic dataset (Fig. 2), were mutated to alanine (A) to mimic a dephosphorylated state or to aspartic acid (D) to mimic constitutive phosphorylation. When expressed in iNGN2 neurons and challenged with Tau seeds, the phospho-dead Tau mutant at GSK3 sites (T217A/T231A/S396A) displayed markedly reduced aggregation compared with wild-type Tau (Fig. 8D–F). In contrast, the phospho-mimetic Tau mutant at GSK3 sites (T217D/T231D/S396D) significantly enhanced aggregation, underscoring the direct role of phosphorylation at these sites in driving Tau pathology. The MARK2 site mutations yielded a distinct phenotype: both phospho-mimetic (S262D/T356D) and phospho-dead (S262A/T356A) mutants produced a robust increase in aggregation, even in the absence of exogenous seeding (Fig. 8D, F), suggesting these sites are also important regulators of aggregation, with different mutations potentially impacting microtubule binding. Importantly, mutations at the canonical AT8 epitope (S202/T205) did not alter aggregation, suggesting that phosphorylation at these sites is a downstream marker rather than a driver of the aggregation process. Together, these findings provide strong evidence that phosphorylation at GSK3 and MARK2 target sites is a critical prerequisite for Tau aggregation, whereas phosphorylation at AT8 sites is dispensable. Consistent with and extending prior studies,^40,41,42^ our results establish a mechanistic framework in which specific kinase-driven modifications directly initiate Tau misfolding.

**Figure 8.**
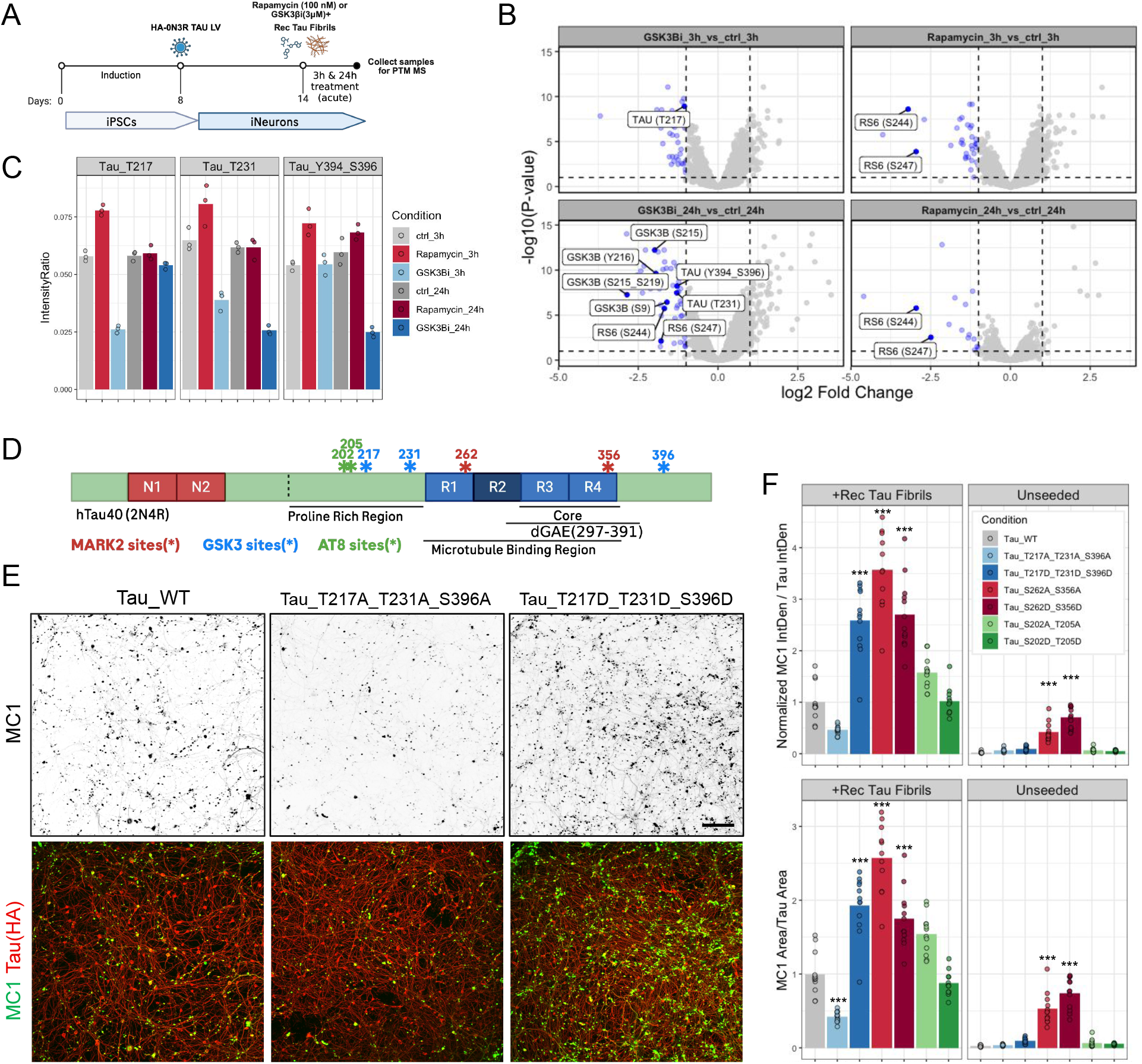
Tau Aggregate Formation Depends on MARK2 & GSK3 Phosphorylation Sites. **(A)** Schematic representation of cell lysate preparation for phosphoproteomic analysis following 3-hour and 24-hour treatments with 100 nM rapamycin (mTOR inhibitor) and 3 µM AZD1080 (GSK3 inhibitor) in the Tau OE Aggregation Model. **(B)** Volcano plot showing relative changes (log₂ fold) in phosphopeptide abundances comparing AZD1080- and Rapamycin-treated cells to vehicle control. Highlighted are phosphorylation changes on Tau, GSK3, and the mTOR target RS6. **(C)** Quantification of selected Tau phosphopeptide intensities showing significant changes following AZD1080 treatment from samples described in (A). **(D)** Schematic representation of Tau phosphorylation sites (*) mutated in this study. Phosphorylation sites targeted by GSK3B are shown in blue, those targeted by MARK2 in red, and AT8 epitope sites in green. All mutations were introduced in the context of the 2N4R Tau isoform (hTau40). **(E)** Representative immunofluorescence images of iNGN2 neurons transduced with wild-type Tau (WT) or Tau mutants T217A_T231A_S396A and T217D_T231D_S396D (mutants S262A_S356A, S262D_S356D, S202A_T205A, and S202D_T205D not shown), seeded or unseeded with recombinant Tau fibrils for 14 days, then stained with MC1 (green) or HA (red, OE Tau) antibodies. Scale bar, 200 µm. **(F)** Quantification of Tau aggregation efficacy as measured by MC1 Integrated Density/Tau Integrated Density (upper panel) and MC1 Area/Tau Area (lower panel) from neurons described in (E). Phosphorylation sites targeted by GSK3B are represented in blue bars, MARK2 sites in red bars, and AT8 sites in green bars; n = 11-12. One-way ANOVA was performed within each seeding condition, followed by Tukey’s post-hoc test. Significance is shown relative to Tau_WT. *p < 0.05, **p < 0.01, ***p < 0.001.

## DISCUSSION

Aggregation and deposition of insoluble Tau protein is a defining pathological hallmark of AD and other tauopathies. In AD, Tau pathology develops in a stereotyped manner across anatomically connected brain regions, suggesting trans-synaptic propagation. The regional burden of Tau pathology correlates closely with clinical decline, emphasizing its central role in disease pathogenesis. Modeling Tau aggregation and pathology in preclinical systems has been challenging, particularly for wild-type Tau, which resists aggregation under physiological conditions.

Here, we describe two reproducible human iPSC-derived neuron models that recapitulate key molecular and cellular features of AD tauopathy. In the seeding model, recombinant disease-relevant pre-formed Tau fibrils (PFFs) are internalized by iNGN2 neurons, leading to progressive, time-dependent intracellular aggregation of endogenous wild-type Tau. In the Tau overexpression model, aggregation is accelerated, enabling biochemical characterization and small-molecule screening. Both models rely exclusively on wild-type Tau, circumventing the need for aggregation-prone mutant forms of Tau commonly used for modeling tauopathy (eg. P301L, P301S). While such mutations can facilitate aggregation formation, they are known to form distinct fibril structures and may bypass some events, such as early post-translational modifications, required to drive aggregation of the wild-type protein. Together, these systems reproduce hallmark events of tauopathy, from initial seeding to synaptic dysfunction, within a human neuronal context suitable for mechanistic investigations into the aggregation process and cellular responses to it.

### Fibril Structure and Subcellular Distribution

Biochemical isolation of insoluble fractions and cryo-ET imaging confirmed the formation of mature fibrillar Tau in our models. To our knowledge, this represents the first demonstration of Tau PHF formation by cryo-ET directly within the native neuronal cell environment. Notably, our systems captured fold- and strain-specific features of Tau aggregation. Recombinant Tau seeds, which are structurally heterogeneous, produced diverse filament structures, including PHFs, whereas AD-derived Tau seeds generated morphologically homogeneous PHFs that closely resemble those observed in human brain tissue, suggesting that features of the initiating seed can be retained during intracellular fibril propagation. Recently reported methods for generating recombinant PFFs with more reproducible AD-like PHF structure may increase the fidelity of recombinant seeding material.^43^

Cryo-ET further revealed that fibrils were distributed throughout the cytoplasm and not encapsulated within vesicular compartments, consistent with reports that Tau fibrils escape the endolysosomal compartments.^44^ Importantly, Tau fibrils accumulated at synapses and physically entrapped synaptic vesicles, correlating with disrupted network activity in seeded cultures. These findings establish a direct mechanistic link between Tau aggregation to synaptic dysfunction, suggesting that early synaptic impairment is a proximal consequence of intracellular Tau pathology.

### Phosphorylation-Dependent Mechanisms of Aggregation

Hyperphosphorylation of Tau is tightly associated with aggregation and neurodegeneration. In our model, seeded cells displayed abundant Tau phosphorylation correlating with aggregation. By immunocytochemistry, we confirmed that Tau aggregates were heavily phosphorylated at multiple disease-associated sites, including Ser202/Thr205 (AT8), Ser409 (PG5), Thr231 (AT180), Ser422, and Thr217. Global phosphoproteomic analysis revealed additional Tau phosphorylation events in total cell lysates, some of which appear to be critical for aggregation. Integrated phosphoproteomic profiling and pharmacological perturbations identified MARK2 and GSK3 as central mediators, with phosphorylation at S262, T356, T217, T231, and S396 emerging as key triggers of aggregation. While aggregated Tau is hyper-phosphorylated at many residues, the contribution of specific phosphorylation events to Tau aggregation has not been demonstrated. Using site-directed mutagenesis, we provide direct causal evidence that modifications at the S262/T356, and T217/T231/S396 sites actively promote misfolding, whereas AT8 sites (S202/T205) represent downstream consequences rather than initiating events.

These phosphosites cluster within the proline-rich, microtubule-binding and C-terminal regions of Tau, where phosphorylation may disrupt microtubule interactions and/or destabilize the “paperclip” conformation of soluble tau, exposing aggregation-prone motifs.^45^ The relevance of these sites is further supported by human postmortem and clinical studies implicating pS262 and pT356 as candidate biomarkers of early soluble assemblies,^46^ and the strong performance of T217 and T231 as robust AD biomarkers.^47^ Together, our findings define a phosphorylation-dependent mechanism in which early modification at microtubule-binding domains and surrounding GSK3 motifs primes Tau for pathogenic conversion.

### Signaling Pathways Regulating Tau Aggregation

Pharmacological screening identified a number of signaling pathways that can modulate Tau aggregation in human neurons. Across our models, we consistently identified the PI3K/mTOR/GSK3 axis as a key regulatory pathway. Compounds that reduced aggregate burden also restored synaptic clustering, indicating that attenuating Tau aggregation can directly improve synaptic integrity. FRET-based seeding assays revealed that PI3K inhibitors and, to a lesser extent, mTOR inhibitors act at early stages of the aggregation cascade, reducing fibril uptake and seeding efficiency rather than merely modulating Tau expression or phosphorylation. This observation is consistent with the established role of PI3K in endocytosis and vesicle trafficking.^48^ In contrast, GSK3 inhibition predominantly reduced phosphorylation at aggregation-driving sites without affecting initial seeding events. These data integrate well with prior studies connecting PI3K–AKT–mTOR and GSK3 signaling to Tau pathology in AD.^49,50^

### Comparison with Existing iPSC Models

Several iPSC-derived neuronal tauopathy models have been reported in the literature.^51,52,53,54,55,56,57,22,58,59,60,61,62^ Some of these models demonstrated robust Tau aggregation.^22,57^ Most required pathogenic Tau mutations (e.g., P301L, A152T) in combination with seeding to drive aggregation. Mutations in the Tau protein drive conformational changes that limit the types of biochemical pathways and molecular interactions it can participate in. To more accurately model AD, which lacks Tau mutations,^63^ a model featuring wild-type Tau is desirable. A wild-type Tau aggregation model seeded with AD Tau was recently described^22^. While patient-derived seeds are unparalleled for generating physiologically accurate PHFs, they are limited in availability, vary in seeding potency across individuals and brain regions, and can persist extracellularly in cultures, complicating intracellular aggregate detection by immunofluorescence.

Compared with prior approaches, our model employs wild-type Tau and recombinant seeds that are rigorously characterized. Recombinant dGAE (297–391) fibrils, which are MC1- and phospho-Tau–negative potently trigger aggregation of wild-type 0N3R Tau. NGN2-based differentiation yields a homogeneous excitatory neuron population, enabling robust and reproducible aggregation across replicates. The combination of endogenous and overexpression models allowed both mechanistic studies and a medium-throughput automated screen of an 800 compound library - the first, to our knowledge, conducted in human iPSC-derived neurons using wild-type Tau aggregation as a readout.

### Therapeutic Context and Implications

Despite extensive efforts, clinical trials targeting Tau have produced limited success to date.^64^ Multiple passive immunotherapies directed against extracellular Tau have been evaluated clinically.^65,66,67,68,69^ While several achieved some level of target engagement of soluble extracellular Tau in cerebrospinal fluid (CSF), there has been minimal effect on Tau PET or cognition,^66,70^ suggesting that targeting extracellular Tau is insufficient to provide meaningful clinical benefit. Targeting the intracellular pool of Tau has also proven challenging. Small molecule approaches, including aggregation inhibitors,^71^ microtubule stabilizers,^72^ GSK-3 kinase inhibitors,^73^ mTOR inhibitor rapamycin^74^ and O-GlcNAcase inhibitors,^75,76,77^ showed little or no biomarker or clinical benefit. However, definitively assessing the therapeutic efficacy of these compounds is difficult, as these studies generally failed to demonstrate target specificity and engagement within the brain. In contrast, oligonucleotide-based therapies that act to reduce new Tau protein synthesis, such as the antisense oligonucleotide (ASO) BIIB080, demonstrate robust target engagement, with reductions of soluble CSF Tau and Tau PET signal in AD patients,^78^ with early indications of potential clinical benefit. However, the requirement for repeated intrathecal administration may limit long-term treatment for many patients.

Our data extend the mechanistic basis for new therapeutic strategies by identifying critical steps in the Tau aggregation process that are amenable to intervention. Specifically, Tau fibril uptake and seeding in our neurons is dependent on PI3K signaling. Furthermore, we identified key phosphorylation sites on Tau that appear to act as molecular switches for aggregation. Some of these sites are associated with early Tau assemblies,^46^ an ideal target for intervention. Interfering with these processes could slow down or block Tau aggregation at different steps, and potentially provide clinical benefit to AD patients. PI3K, GSK3 and MARK2 are broad-acting kinases, and their therapeutic use in AD may be limited by their regulation of other substrates and biological processes. Physicologically relevant cell models such as the ones presented here provide a platform for further discovery of selective effectors or downstream modifiers that can attenuate or reverse Tau pathology in neurons.

### Limitations and Future Directions

While our models capture key aspects of tauopathy, several limitations remain. First, recombinant fibrils used for seeding differ from the PHF-dominated aggregates observed in AD brain tissue; developing methods to generate AD-like fibrils more reproducibly will improve disease relevance. Whether fibril structure itself dictates pathology, or instead reflects the site of initiation, remains an open question. Second, NGN2 neurons constitute a homogeneous excitatory population, lacking the cellular diversity of the human brain. Incorporating astrocytes, microglia, and oligodendrocytes will be essential to better recapitulate the multicellular environment that shapes Tau pathology. Third, while we identified kinase pathways and specific phosphorylation sites that regulate aggregation, future studies should systematically dissect the functional consequences of individual modifications observed in our PTM analyses. Notably, some residues such as pS305 have been reported to inhibit aggregation, underscoring the complexity of phosphorylation events in Tau pathology.^79^ Our data also implicate Chk1 as a potential regulator that may drive aggregation by increasing phosphorylation at sites such as S262 and T356—consistent with prior findings in Drosophila models^80^ and *in vitro* studies.^81^ Defining Chk1’s role in Tau regulation will be an important avenue for future work. Finally, extending these findings to *in vivo* models that reproduce relevant Tau fibril confirmations and disease progression will be critical for evaluating therapeutic potential.

## ACKNOWLEDGMENTS

We thank members of the Genentech Neuroscience Department for their helpful suggestions during the study. We are especially grateful to Sofia Lövestam for her advice on generating the dGAE Tau fibril preparation, and to Nicole Koutsodendris for isolating the AD Tau seeding material. We thank Christine Tam and the BioMolecular Resources (BMR) group at Genentech for their support in generating DNA constructs. Finally, we thank Ada Ndoja and Michelle Chen for their critical review of the manuscript. All studies were funded by Genentech, Inc., South San Francisco, CA.

## AUTHOR CONTRIBUTIONS

J.L. designed and performed experiments, coordinated the study, analyzed data, and wrote the manuscript. X.S. designed and performed experiments in the non-OE Tau model and analyzed data. L.P. and A.O. designed, performed, and analyzed proteomic experiments. P.D. designed, performed, and analyzed cryo-ET experiments. T.W. generated recombinant Tau fibril material. M.J. designed, performed, and analyzed CryoEM experiments. Q.Z. and Z.T. designed, performed, and analyzed the HTS experiments. M.A. and A.B. analyzed imaging experiments. D.S. performed protein purification, M.C. contributed to MS data analysis, J.M. designed the compound library. C.C.H., C.J., C.E., Z.T., A.R., and A.O. supervised the research. J.A. supervised the research and wrote the manuscript. All authors read and edited the manuscript.

## RESOURCE AVAILABILITY

### Lead contact

Requests for further information and resources should be directed to the lead contact, Dr. Jasi Atwal.

### Materials and data availability

Requests for further information, resources, or reagents should be directed to the lead contact and will be fulfilled under a material transfer agreement. Any additional information required to reanalyze the data reported in this paper is also available from the lead contact upon request.

## MATERIALS & METHODS

### Recombinant Tau Fibril Preparation

*In vitro* assembly of monomeric, recombinant Tau into a mixture of filaments adopting the AD proto-filament fold (i.e. Paired Helical Filaments (PHF), Triple Helical Filaments (THF) and Quadruple Helical Filaments (QHF)) was adapted with modifications from a protocol previously reported^24^. For protein expression, a DNA sequence encoding a recombinant fragment of human MAPT (Uniprot ID P10636-8, residues 297-391, frequently termed dGAE Tau) was cloned into a modified pET-52b vector and transformed into *E. coli* BL21 (DE3) competent cells. 50 ug/mL carbenicillin was supplemented during all subsequent steps of protein expression. The transformation reaction was plated onto LB agar and a single transformant colony was inoculated into a LB media starter culture and incubated overnight at 37 °C. Starter culture was used to inoculate 1 L TB autoinduction media cultures grown in UltraYield flasks (Thomson, 931136B) at 37 °C for 8 h, followed by 16 h at 24 °C. Cells were pelleted by centrifugation and frozen.

For recombinant Tau purification, cell pellets were thawed and resuspended in [50 mM MES, pH 6.3, 10 mM EDTA, 10 mM DTT], and supplemented with cOmplete EDTA-free Protease Inhibitor Cocktail (Roche). Resuspended cells were lysed by sonication [Branson Digital Sonifier SFX 550, 40% amplitude for 7 minutes, 5s on/10s off]. Lysed cells were centrifuged at 20,000 x g for 35 min at 4 °C, the clarified lysate was filtered (0.45 um cutoff) and loaded onto a HiTrap CaptoS 5 mL cation exchange column (Cytiva, 17544123). The column was washed with 10 column volumes (CV) of Wash Buffer [50 mM MES pH 6.0, 10 mM EDTA, 10 mM DTT], and Tau protein was eluted with a 20 CV linear gradient to Wash Buffer + 1 M NaCl. Fractions containing recombinant Tau were pooled and precipitated by adding 0.3 g/mL of ammonium sulfate and rotating the mixture for 30 min at 4 °C. Precipitated Tau was pelleted by centrifugation (20,000 x g, 35 min, 4 °C), the pellet was resuspended in 10 mM Sodium Phosphate pH 7.2, 10 mM DTT, and polished by gel filtration chromatography (HiLoad 16/600 Superdex 75 pg, Cytiva, 28989333), typically yielding ∼15 mg Tau protein per 1 L *E.coli* cell culture. Peak fractions were concentrated to 20 mg/mL (based on A_280_), flash frozen in 40 µL aliquots and stored at -80 °C.

For filament assembly, an input monomeric Tau mixture was prepared by thawing recombinant Tau and diluting to 8 mg/mL into a final buffer condition of [10 mM Sodium Phosphate pH 7.2, 100 mM MgCl_2_, 10 mM DTT, 2 uM Thioflavin T (ThT)]. For generation of large batches of recombinant Tau fibrils, this Tau mixture was dispensed at 40 µL per well across multiple wells of a Corning 384-well Black/ Clear Flat bottom plate (Corning 3544), with a typical scale of 40 reaction wells per experiment. In-plate filament assembly was carried out using a FLUOstar Omega Plate-reader (BMG Labtech) incubated at 37 °C while applying a double-orbital shaking pattern at 700 rpm for 16 h. Filament formation reactions proved highly variable well-to-well as tracked via increase in ThT fluorescence. Filaments generally formed after 4-8 h, with reactions producing either a distinct high or low endpoint ThT fluorescence signal (∼5x signal difference). Characterization by cryoEM revealed high ThT endpoint reactions contained higher proportions of PHFs, THFs and QHFs, whereas low ThT endpoint reactions predominantly contained flat filaments. High endpoint ThT reactions were pooled into large batches, which were further characterized by cryoEM for filament structure, and for seeding potential in HEK293 Tau biosensor and iNGN2 neurons.

### AD Tau Fibril Preparation

All reagents were prepared using deionized water. The Extraction Buffer consisted of 10 mM Tris-HCl (pH 7.5), 10% sucrose, 0.8 M NaCl, 5 mM EDTA, 1 mM EGTA, and protease/phosphatase inhibitors. The Resuspension Buffer contained 10 mM Tris-HCl (pH 7.5) and 100 mM NaCl, along with protease/phosphatase inhibitors. A 30% stock of N-lauroylsarcosinate (sarkosyl) was used for detergent solubilization.

Approximately 10 g of human tissue was homogenized in 100 mL of ice-cold Extraction Buffer using a Polytron homogenizer. The homogenate was then centrifuged at 20,000 g for 20 minutes at 4°C. The supernatant was collected, and the pellet was re-homogenized in fresh Extraction Buffer and centrifuged again under the same conditions. This second supernatant was combined with the first one.

The combined supernatant was treated with a final of 1% sarkosyl and incubated at room temperature for 1 hour with stirring. This mixture was then subjected to ultracentrifugation at 100,000 g for 1 hour at 4°C. The resulting pellet was resuspended in Resuspension Buffer and diluted 30-fold with Extraction Buffer, re-homogenized, and centrifuged at 20,000 g for 20 minutes. The resulting supernatant underwent a final ultracentrifugation at 100,000 g for 1 hour. The final pellet was resuspended in 250 µL of Resuspension Buffer for further analysis.

### CryoEM of Tau Fibrils

For all cryoEM experiments, 2.5 microliters of sample were applied to glow discharged amorphous Ni-Ti holey foil grids (Molecular Dimensions Inc., Holland, OH, USA), and then plunge frozen using a Vitrobot (Thermo Fisher Scientific, Inc., Waltham, MA, USA). Grids were screened for quality and imaged for structure determination using a Titan Krios TEM equipped with a Falcon 4i direct detector and Selectris energy filter. Images were collected for both screening and structure determination at a nominal magnification of 81kx (1.467 Å/px), and a total electron fluence of 40 e/Å^2. For screening datasets, image processing was performed using CryoSPARC.^82^ Following movie alignment and CTF estimation, filaments were traced using the *ab initio* filament tracer tool and segments were extracted using a box size of 600 Å and 14.7 Å repeat distance. Filament segments were sorted via 2D classification and segment classes were manually binned into three categories: PHF/THF/QHF, twisting but non-PHF fold, and nontwisting “flat” filaments. Datasets used for structure determination were processed using Relion, according to previously established protocols.^83,84^

### Culturing of iPSCs and NGN2 Neurons

Cell culture protocols were modified from a previously published protocol.^85^

All cells were maintained in 37°C 5% CO2 incubators unless otherwise indicated.

iPSCs (iP11N) were maintained in mTeSR Plus medium on tissue culture plates coated with iMatrix-511 (Takara) in 37°C 5% CO_2_ incubators. Cells were passaged as clumps with ReLeSR every 3-4 days at a ratio of 1:6. Cells were checked every 5 passages for mycoplasma contamination using MycoAlert detection kit and genetic stability was confirmed with Human Pluripotent Stem Cell Genetic Analysis Kit.

On day 0 of neuron induction, iPSCs were dissociated with Accutase and plated at densities 50,000/cm^2^ on iMatrix-511-coated plates in mTeSR Plus supplemented with RevitaCell (1x Invitrogen) and doxycycline (3 µg/ml). Then, medium was changed into DMEM/F12 supplemented with Glutamax (1x), NEAA (1x), B27 without VA (1x), N2 Max (1x), doxycycline (3 µg/ml), DAPT (10 μM), SB431543 (10 μM), Noggin (100 ng/mL), and XAV939 (2 μM). Daily media changes were performed on days 1-4. On day 5, medium was changed into Neurobasal supplemented with Glutamax (1x), B27 with VA (1x), dcAMP (500 µM), Ascorbic Acid (200 µM), BDNF (20 ng/mL), GDNF (20 ng/mL), DAPT (10 μM), doxycycline (2 µg/ml), and Arabinocytidine (5 μM). On Day 7, cells were dissociated with Papain and DNase. Cells were cryopreserved with CryoStor CS10 and CoolBoxes in -80°C, and then stored in liquid nitrogen till use.

Cryopreserved day 7 neurons were thawed in a 37°C water bath and plated on tissue culture plates coated with PDL and iMatrix-511. Neuronal maturation media contains Neurobasal supplemented with Glutamax (1x), B27 with VA (1x), dcAMP (500 µM), Ascorbic Acid (200 µM), BDNF (20 ng/mL), GDNF (20 ng/mL). Mitomycin C (0.25 μM) was included in the plating media to remove reminiscent progenitors. Cells were maintained in 37°C, 5% CO_2_ incubators with weekly partial media changes.

### Seeding of iNGN2 Neurons

Cryopreserved day 7 neurons were thawed in a 37 °C water bath and plated on tissue culture plates as described above. For most experiments a total of 15,000 live cells per well were plated into 384-well Poly-D-lysine–coated PhenoPlates (Revvity, Cat# 6057500), or 40,000 live cells were plated into 96-well Poly-D-lysine–coated PhenoPlates (Revvity, Cat# 6055500).

On day 8, cells were transduced with an HA-0N3R Tau or Tau Phosphomutant (Figure 8) containing lentivirus (see Key Resources Table) at an MOI of 3–5, which resulted in >85% of cells being HA-Tau positive at the time of detection (data not shown). On day 15, cells were treated with 2 µg/mL recombinant Tau fibrils. Cultures were maintained in 37°C, 5% CO₂ incubators with weekly partial medium changes until fixation in 4% paraformaldehyde (PFA, Electron Microscopy Sciences, cat#15710) for the period specified in the corresponding figures/figure legends. In the non-OE seeding model, no lentivirus was added.

### Immunofluorescence (IF)

Immunostaining was performed following our previously published protocol^86^ with minor modifications. Fixed cells were permeabilized in 0.5% Triton X-100 in PBS for 5 min, followed by blocking for 1 h at room temperature in buffer containing either 3% bovine serum albumin (BSA), 2.5% normal donkey serum (NDS), and/or 10% normal goat serum (NGS). Cells were then washed three times with PBS and incubated overnight at 4 °C with primary antibodies diluted in blocking buffer (3% BSA, 2.5% NDS, or 10% NGS). Following three additional PBS washes, cells were incubated for 1h at room temperature with secondary antibodies in blocking buffer (3% BSA, 2.5% NDS, and/or 10% NGS) together with Hoechst 33342 (1:1000). All antibodies are listed in the Key Resources Table.

### Tau aggregate imaging and analysis

High-throughput imaging was performed on an ImageXpress Confocal HT.ai High-Content Imaging System (Molecular Devices) equipped with an LDI illuminator (89 North) (Fig. 1C, D, F; Fig. 4A; Fig. 5–6; Fig. 8; Supplementary Fig 2). Images were acquired using 10× Plan Apo 0.45 NA, 20× APO LWD 0.95 NA WI, or 40× APO LWD 1.15 NA WI (Nikon) objectives with appropriate filters and excitation lines for either secondary antibodies or expressed fluorescent proteins. Image analysis was carried out in MetaXpress (v6.7.2.290, Molecular Devices) using the Custom Module Editor. For Tau-biosensor–expressing HEK cells, nuclei were identified by DAPI staining, and expressing cells were defined by low YFP thresholding after Gaussian blurring (sigma = 5). Tau aggregates were segmented by applying a high adaptive YFP threshold empirically defined for each experiment to identify bright objects between 0.5 and 10 µm in diameter. In NGN2 neurons, Map2- or Tubb3-positive areas were quantified using adaptive thresholds 500–1000 AU above local background, while AT8-positive regions were segmented at 1000–2000 AU above local background with widths between 0.1 and 5 µm. Similarly, p62- or synaptophysin-positive areas and puncta counts were quantified using adaptive thresholding.

### High Throughput Automation and Screening of NGN2 Neurons

A library of target-annotated compounds was assembled from compounds synthesized at Genentech or purchased from MedChemExpress. Literature-derived annotations for targets were manually reviewed and modified as necessary. Compounds were dissolved in DMSO as 10mM stock solutions and dispensed into 384-well Low Dead Volume plates (Beckman Coulter).

Cryopreserved day 8 human iP11N-derived neurons were thawed in a 37°C water bath on day 0, and viable cells were counted. Using an Agilent Bravo liquid handling system with custom methods, 15,000 cells per well were dispensed into 384-well Poly-D-lysine–coated PhenoPlates (Revvity, Cat# 6057500) pre-coated with iMatrix-511 (Takarabio Cat # T303).

On day 1, Tau ON3R lentivirus (multiplicity of infection [MOI] = 3) was added to induce ON3R Tau overexpression in NGN2 neurons for one week (details of virus generation provided in a separate section). Compounds were acoustically transferred from stock plates into intermediate plates using a Beckman Coulter Echo liquid handler. On day 7, culture medium was aspirated and replaced with fresh medium supplemented with Tau recombinant fibrils (2 µg/mL) using an Agilent BioTek EL406 washer. Medium changes were performed on days 7, 12, and 17. At each medium change, gProbe compounds were re-dispensed into the cell plates using the Agilent Bravo system to achieve the screening concentration, e.g. 3 uM for the primary screen.On day 21, cells were fixed with 4% paraformaldehyde and immunostained as described above.

All liquid handling steps (aspiration and dispensing) were performed using the BioTek EL406 paired with a BioStack loader and custom protocols. Confocal imaging was conducted on a Molecular Devices ImageXpress system, and image analysis was performed using Molecular Devices MetaXpress (Molecular Devices) software as described above.

### Quantification and Statistical Analysis of the Screen Data

#### Normalization

Toxicity (MAP2 area) was normalized to the mean of on-plate DMSO controls and expressed as a percentage. Efficacy (AT8 immunostaining) was calculated as integrated AT8 area per MAP2+ nuclei count, normalized to plate-matched DMSO controls and expressed as a percentage.

#### Toxicity filtering

treatments (defined as the combination of compound and dose) with <60% average MAP2 area relative to DMSO controls were excluded from further analysis.

#### Statistical testing

For each treatment group, a two-tailed Student’s *t*-test was performed against DMSO controls from the same plate. P-values were -log10 transformed for visualization. False discovery rate (FDR) correction (Benjamini–Hochberg) was used to adjust p-values for multiple testing.

#### Effect size estimation

The strictly standardized mean difference (SSMD) was calculated for each treatment versus its DMSO controls, providing a measure of effect size.

#### Hit definition

Compounds were classified as reducers if SSMD < –2 and *p* < 0.01, and as boosters if SSMD > 2 and *p* < 0.01. Non-significant treatments were labeled as non-hits.

### Tau Hibit-Halo and Tau Knockout (KO) iPSC Cell Line Generation

iPSC engineering method was modified from a previously published protocol.^85^

On day 0, one well of a 24 well plate was coated with GelTrex for 1h at 37°C. After coating, GelTrex was removed and 2 mL media was added to the well and returned to the incubator for temperature and gas equilibration. For engineering the Hibit-Halo line, the media contains mTeSR Plus supplemented with 1x CloneR2 and 1x HDR Enhancer V2. For the MAPT KO line, the media contains mTeSR Plus supplemented with 1x CloneR2.

RNP (Ribonucleoprotein) was assembled by incubating 500 pmol of gRNA with 2 μL of HiFi Cas9 nuclease V3 for 30 min. iPSCs at 75% confluency were dissociated using Accutase. 1.5 million cells were pelleted by centrifugation at 300g for 5min. Cells were resuspended with 20ul P3 solution and mixed with RNP before transferred to one well of a nucleofection strip (Lonza). In the case of Hibit-Halo tags KI line, single stranded donor DNA (200 pmol) were added to the nucleofection mix. Nucleofections were performed with program CM113. After nucleofection, cells were transferred to the pre-equilibrated well in the 24 well plate and returned to the incubator. For the Hibit-Halo line, cells were kept in a 32°C incubator for 2 days to increase homology directed repair rate. For the MAPT KO line, cells were kept in a 37°C incubator.

Media was changed into mTeSR Plus on day 1. On day 2, cells were dissociated into single cells with Accutase and 25,000 cells were plated on 10cm dishes coated with GelTrex in mTeSR Plus supplemented with ClonR2. ClonR2 were removed 48h later and iPSCs were maintained for 1 week. 96 colonies were picked from the plate under an EVOS microscope in a biosafety cabinet. Half of each colony were plated into a 96 well plate coated with Geltrex in mTeSR Plus, and the genomic DNA of the other half were extracted with QuickExtract. Genotyping was performed by PCR with Q5 Hot Start High-Fidelity 2X Master Mix at an annealing temperature of 60°C. PCR products containing correct bands were further sequenced by Sanger sequencing. Positive clones were further expanded from the 96 well plate to 6 well plates. Candidate clones were QC’d using Karyostat Plus, MycoAlert, and neuronal induction. Pluripotency markers OCT3/4, NANOG, and SOX2 were checked with immunofluorescence. Clones passing QC were further expanded on iMatrix and banked with CryoStor CS10.

### Tau level measurements (HaloTag-based readout)

Bright-field images of Hibit-Halo-tagged iNGN2 neurons treated with compounds in 384 well plates were acquired on Cytation 5 high-content imager for live cell counting. Cells were then lysed with Nano-Glo® HiBiT Lytic Detection kit (20 µL/well; Catalog Number: N3030; Promega). Lysed cells were incubated at 22°C for 20 min. Luminescence readings were performed with a Synergy Neo2 plate reader.

### CRISPR KO of NGN2 Neurons

For CRISPR–Cas9 RNP electroporation, ribonucleoprotein (RNP) complexes were assembled by incubating 1 µL SpCas9 Protein V3 (500 µM; IDT, cat# 1081059), 1 µL Duplex Buffer (IDT, cat# 11010301), and 3 µL gRNA (Synthego; Gene Knockout Kit, 3 guides per gene, 500 pmol) for 15 min at room temperature. Approximately 0.5 × 10⁶ cells (Day 7) were pelleted at 400 × g for 5 min and resuspended in 20 µL P3 Buffer (Lonza). For electroporation, 18 µL of the cell suspension was mixed with 4 µL of RNPs and transferred into one well of a nucleofection strip (Lonza 4D) using the CV-110 program. Following electroporation, 30,000 cells per well were plated into 384-well assay plates in neuronal medium supplemented with 10 µM Y-27632 (Stem Cell Technologies, cat# 72302) to enhance neuronal survival.

### FRET-Based Tau Uptake Sensor in NGN2 Neurons

Tau KO neurons were plated on Day 7 at a density of 15,000 cells per well in 384-well plates. On day 8, cells were transduced with mClover3- and mRuby3-tagged Tau^P301S^RD-carrying lentiviruses at an MOI of 4 (see Key Resources Table). One week later, cultures were treated with Tau fibrils (2 µg/mL; StressMarq, cat# SPR-480) for 3 days in the presence of compounds at 3 µM. Following the 3-day treatment, FRET (Fluorescence Resonance Energy Transfer) signal was measured.

### FRET imaging and analysis of NGN2 Neurons

FRET measurements of mClover3- and mRuby3-tagged TauP310SRD in iNGN2 neurons were acquired on a Leica Thunder DMi8S inverted microscope (Leica Microsystems) with motorized stage and AutoFocus Control. Samples were illuminated by a Spectra X light engine (Lumencor) using 438 and 555 nm excitation lines through a CFP/YFP dichroic, a HC PL APO 20x NA 0.80 air objective (Leica Microssystems) and 510 and 590 nm emission filters and imaged onto a K8 sCMOS camera (Leica Microsystems). Images were analyzed with Cellprofiler v4.2.5^87^ To measure FRET-positive areas, we thresholded the FRET signal using adaptive thresholds and the “Robust Background” algorithm looking for signals higher than three standard deviations of the mean image intensity.

### Sarkosyl Fractionations and Western blotting

Cells were gently washed with PBS and lifted from the wells by adding ReLeSR™ Solution (Catalog # 100-0483; Stem Cell Technologies). Subsequently cells were spun down at 500xg for 10 minutes and Extraction buffer (800 mM NaCl, 20 mM Tris-HCl pH 7.4, 5 mM EDTA, 15% sucrose, 1% sarkosyl, protease inhibitor (cat# 4693132001, Roche)) was added to cell pellets. Samples were sonicated in the cold room using a metal probe ultrasonicator for 15 s at setting 4.5 (11–12 W) and incubated at 37°C with shaking at 300 RPM for 30 min. Lysates were centrifuged at 50,000 RPM for 45 min at 22°C, and the resulting supernatant was collected and retained (sarkosyl soluble fraction), while the pellet was resuspended in 50 μL of buffer containing 50 mM Tris and 150 mM NaCl. The suspension was clarified by centrifugation at 3000 × g for 5 min, and the supernatant was transferred to a new tube. This supernatant was centrifuged again at 50,000 RPM for 45 min at 22°C, after which the final pellet was resuspended in 15 μL of buffer (50 mM Tris, 150 mM NaCl) and subjected to a final round of sonication in a water bath sonicator for 45 s (15 s on, 10 s off cycles).

For immunoblotting, samples were resolved by SDS-PAGE using precast gels (Invitrogen) and transferred onto nitrocellulose membranes (Bio-Rad). Membranes were blocked with 5% bovine serum albumin in 1× TBST (10 mM Tris-HCl, pH 8.0, 150 mM NaCl, 0.1% Tween-20) and incubated overnight at 4°C with primary antibodies diluted in blocking buffer. Eluted proteins were probed with antibodies (Key Resources Table), each used at 1:1000 dilution. After three 10 min washes in TBST at room temperature, membranes were incubated with IRDye® 800CW donkey anti-rabbit IgG or IRDye® 680 RD donkey anti-mouse IgG secondary antibodies (Li-Cor) and imaged on the Odyssey DLx imaging system (Li-Cor).

### MS Sample Preparation for Tau Seeding and Compound Screening

Samples were lysed in RIPA buffer (Thermo Fisher Scientific, cat# 89901) containing protease and phosphatase inhibitors (Roche). To each 50 µg of protein lysate, sodium dodecyl sulfate (SDS) was added to a final concentration of 5%. Lysates were sonicated using a PIXUL (ActiveMotif), reduced (5 mM dithiothreitol, 45 min at 37°C), alkylated (10 mM iodoacetamide, 15 min at room temperature in the dark), and processed using the S-Trap 96-well Mini plate (Protifi) according to the manufacturer’s protocol. Enzymatic digestion was performed on-plate for (1 hr at 47°C) using a combination of sequencing grade modified trypsin (Promega) and lysyl-endopeptidase (Wako), both at a 1:15 enzyme to protein ratio. Digested peptides were lyophilized in preparation for TMT labeling.

Peptides were resuspended in 100 µL of 100 mM HEPES pH 8.0 and labeled with 18-plex TMTPro reagents. Each vial of TMT reagent was allowed to thaw for 5 min at room temperature, spun down using a benchtop centrifuge, resuspended in 20 µL of anhydrous ACN, and added to each sample. After 1 hr incubation at room temperature, the reaction was quenched by addition of 5 µL of 5% hydroxylamine for 15 min. Labeled peptides were combined, desalted by C18 Sep-Pak (Waters) solid phase extraction, and dried by vacuum centrifugation.

The TMT-labeled sample was redissolved in 0.15% trifluoracetic acid and subjected to centrifugation at 16,000 x g to remove insoluble material with the supernatant taken forward for offline high pH reversed phase liquid chromatography separation on an Agilent 1100 series HPLC system. Peptides were loaded onto an Agilent Zorbax 300 Extend C-18 analytical column (2.1 × 150 mm and 3.5 μm particle size, Part No: 763750-902). Solvent A consisted of 25 mM Ammonium Formate (pH 9.7), while solvent B was 100% acetonitrile. Peptide fractions were collected in 45 s intervals for a total of 63 min with a linear gradient of 15–60% solvent B. In total 96 fractions were collected and every 25th fraction was combined to form a final set of 24 fractions. Samples were dried in a speed vac and desalted using C18 cartridges (5 µL) on an AssayMAP (Agilent). The eluates were split, with 10% reserved for global protein profiling and the remaining 90% used for phosphoproteomics. Global protein profiling samples were speed vac dried to completion. Phosphoproteomics samples were further concatenated from 24 fractions into 8, combining every 8^th^ sample and then lyophilized prior to phosphoenrichment.

### Phosphopeptide enrichment

Samples were reconstituted in two steps, first by addition of 80 µL 50% ACN/0.1% trifluoracetic acid (TFA) with vortexing for 10 minutes followed by addition of 120 µL of 99.9% ACN/0.1% TFA for a final concentration of 80% ACN. Phosphopeptide enrichment using immobilized metal affinity chromatography was performed on an AssayMAP using Fe(III)-NTA cartridges. Eluates were subsequently desalted using C18 cartridges (5 µL) on the AssayMAP and speed vac dried to completion prior to mass spectrometry analysis.

### Mass Spectrometry: Global Protein Profiling of Tau Seeding

Samples were resuspended in solvent A (2% acetonitrile (ACN)/0.1% formic acid (FA)) and nLC-MS/MS analysis was performed on 10% of each fraction on an Orbitrap Ascend mass spectrometer (ThermoFisher) coupled to a Dionex Ultimate 3000 RSLC nano Proflow system (ThermoFisher) equipped with an Aurora Series 25 cm x 75 um I.D. column (IonOpticks). Low pH reversed-phase separation was performed at 300 nL/min on a 72.9 min two step linear gradient where solvent B (0.1% FA/98%ACN/2% water) was ramped from 4% to 30% over 68 min and then from 30% to 50% over 4.9 min with a total run time of 95 min. For these analyses, a real-time search against a human database was employed using the capabilities built within the XCalibur method editor. For all runs, intact peptides were surveyed in the Orbitrap (250% normalized AGC target, 120,000 resolution) and the top 20 peptides were selected for MS2 fragmentation (150% normalized AGC target; CID, 30 NCE) and analyzed in the ion trap. Synchronous-precursor-selection (SPS) MS3 scans were analyzed in the Orbitrap at 45,000 resolution with the top 8 most intense ions in the MS2 spectrum subjected to HCD fragmentation at a normalized collision energy of 45%, 300% normalized AGC target, and a max injection time of 350 ms. Raw datafiles were deposited into the MassIVE repository with the identifier: MSV000099016 (reviewer login = MSV000099016_reviewer & password = TAU).

### Mass Spectrometry: Global Protein Profiling of Compound Screening

nLC-MS/MS analysis was performed as described above for the Tau seeding experiment with the following exceptions. Six percent of each sample was injected and analyzed on an Oribtrap Eclipse mass spectrometer (ThermoFisher) equipped with FAIMS ProDuo (ThermoFisher) coupled to a Dionex Ultimate 3000 RSLC nano Proflow system (ThermoFisher) equipped with an Aurora Series 25 cm x 75 um I.D. column (IonOpticks). For these analyses, a real-time search against a human database was employed using an in-house instrument API program called InSeqAPI ^88^ ^89^. Data was acquired within each run at two FAIMS CV values of -40V and -60V. The two-stage linear gradient was modified, with buffer B (98% ACN, 2% H2O, 0.1% formic acid) ramping from 4% to 30% over 68 min and then from 30% to 75% over 4.9 min. The top 10 peptides were selected for MS2 fragmentation and synchronous-precursor-selection (SPS) MS3 scans were analyzed in the Orbitrap at 50,000 resolution with the top 8 most intense ions in the MS2 spectrum subjected to HCD fragmentation at a normalized collision energy of 45%, an AGC target of 3.0 x 10^5^, and a max injection time of 400 ms.

### Phosphoprofiling

Phosphoenriched samples were resuspended in solvent A and nLC-MS/MS analysis was performed on 40% of each fraction as described for compound screening global protein profiling with the following modifications. Low pH reversed-phase separation was performed at 300 nL/min on a 150 min two step linear gradient where solvent B (0.1% FA/98%ACN/2% water) was ramped from 4% to 30% over 135 min and then from 30% to 75% over 15 min with a total run time of 185 min. Rather than CID MS2 fragmentation, HCD MS2 (150% normalized AGC target; HCD, 27.5 NCE) was performed.

### Data Processing: Global Protein Profiling

MS/MS spectra were searched using COMET against a concatenated target-decoy database of human proteins (UniProt downloaded January, 2023) containing common contaminant sequences. Search parameters included trypsin cleavage with an allowance of 1 missed cleavage event, a peptide mass tolerance of 20 ppm, and recommended fragment ion settings for low resolution MS/MS with a fragment ion bin tolerance and bin offset of 1.0005 and 0.4, respectively. Searches permitted variable modifications of methionine oxidation (+15.9949 Da) and TMT labeled tyrosine (+304.20715 Da) with static modifications for cysteine carbamidomethylation (+57.0215 Da) and TMT tags on lysine and peptide N termini (+304.20715 Da). A false discovery rate (FDR) of 2% using Percolator was applied at the peptide level to filter peptide spectra matches (PSMs). TMT reporter ions produced by the TMT tags were quantified with an in-house software package known as Mojave by calculating the highest peak within 20 ppm of theoretical reporter mass windows and correcting for isotope purities.^90^

### Data Processing: Phosphoprofiling

Searches were performed as described for Global protein profiling except for missed cleavages which was set to 2 and the addition of phosphorylated serine, threonine, and tyrosine (+79.9663 Da) as allowable variable modifications. Site localization confidence scores for phosphopeptides were calculated using Ascore.

For both Global protein profiling and Phosphoprofiling, the R package MSstatsTMT v2.10.0 was utilized for preprocessing PSM-level quantification to derive protein or PTM quantification, as well as for conducting differential abundance analysis.^91^ MSstatsTMT estimated log2 (fold change) and its associated standard error for each protein or PTM using a linear mixed-effect model. Subsequently, MSstatsPTM v2.6.0 combined the estimated log2 (fold changes) and their standard errors for adjusting protein-level changes in PTM abundance. To test the two-sided null hypothesis of no changes in abundance, the model-based test statistics were compared to the Student t-test distribution with the degrees of freedom appropriate for each protein or PTM. Resulting P values were adjusted to control the FDR with the method by Benjamini-Hochberg.

### Multiple electrode array (MEA) recordings

High density MEA plates were treated and conditioned according to manufacturer directions. Chips were coated with PDL dissolved in Boric acid buffer (250 μg/mL) and iMatrix-511 (1:100 in dPBS). Day 7 neurons were thawed and plated at 0.5 million/chip in a 50 µl droplet of culture media, which contains Neurobasal-A supplemented with Glutamax (1x), B27 with VA (1x), dcAMP (500 µM), Ascorbic Acid (200 µM), BDNF (20 ng/mL), GDNF (20 ng/mL) and mitomycin C (0.25 μM). The next day, wells were flushed with 2 mL of culture media without mitomycin C. The media was changed every 3-4 days.

Neuronal activities were recorded on a Max2 instrument (Maxwell biosystems) under default settings once a week on the same day before media change.

### Grids preparation for cryo-ET

Quantifoil London Finder (H2) gold grids (R2/2, 200-mesh) were prepared a day before plating cells. Grids were first glow discharged (Pelco EasiGlow, 2 min, 15 mA), washed in 70% ethanol, and air-dried before coating in a 1:50 dilution of iMatrix in PBS overnight at 37°C. 200,000 cells were plated in 24-well plastic plates, and 3 grids per well were placed at the bottom of the plate immediately after adding cells. iNGN2 neurons were transduced with an HA-0N3R Tau containing lentivirus at an MOI of 3–5 and treated with 2 µg/mL recombinant Tau fibrils. After 5-6 weeks, cells were plunge-frozen in liquid ethane using back blotting in a Leica GP2 (2 min, 90% humidity, RT).

### Cryo-ET data collection and tomogram reconstruction

Tilt-series were collected on a 300 kV Titan Krios microscope (Thermo Fisher Scientific) equipped with a Falcon 4i direct electron detector and Selectris X Imaging Filter. Multi-frame images were collected using PACE-tomo^92^ script within serialEM^93^ at a pixel size of 1.897 Å (64,000× magnification) or 2.95 Å (42,000× magnification) with a constant defocus of - 6.0 μm and 5 eV energy slit. Dose-symmetric tilt-series were collected from −54° to 54° with 2° increments with a total accumulated dose of ∼120 electrons/Å^2^.

The tilt-series were pre-processed using WarpTools (^94^, https://zenodo.org/records/15749944) for motion correction and CTF estimation. They were subsequently aligned using WarpTools’ Etomo^95^ patch tracking implementation. CTF-corrected tomograms with a pixel size of 15 Å were then reconstructed in WarpTools (https://zenodo.org/records/15749944).

### Tomogram segmentation and visualization

Tomogram segmentation was performed using DragonFly (https://dragonfly.comet.tech). Initially, tomograms were filtered to enhance contrast for the manual annotation of 2D patches. To obtain accurate segmentation of Tau filaments, multiple rounds of model training were required. An initial "universal" model was established and subsequently re-trained on each individual tomogram to refine the segmentation. Segmentations were then exported and visualized in ChimeraX^96^, while tomographic slices were visualized using IMOD.^95^

### HEK Cell Biosensor Assay for Determining Seeding Potential of Tau Seeds

HEK biosensor cells expressing Tau RD-CFP/YFP (kindly provided by Dr. Marc Diamond;^97^) were maintained in DMEM supplemented with pyruvate, D-glucose, and L-glutamine (Thermo Fisher), 10% FBS, and 1% penicillin–streptomycin. One day prior to the seeding experiment, cells were replated into PDL-precoated 96-well plates (Revvity, Cat# 6055500) at a density of 40,000 cells per well and allowed to adhere overnight. For seeding, Tau seed–liposome complexes were prepared by combining Tau fibrils with 3.75 µL Opti-MEM (Gibco) and 1.25 µL Lipofectamine 2000 (Invitrogen), to a final volume of 20 µL per well. Complexes were incubated at room temperature for 30 min prior to addition to the cells. After treatment, cells were incubated for 24 h before analysis.

### Quantification, plotting and statistical analysis

ANOVAs with Tukey’s HSD or Dunnett’s post hoc tests, as well as t-tests, were performed using Prism (GraphPad) or R. Plots were generated with the same software. Multiple images were acquired per well. All *n* values represent well replicates.

## DECLARATION OF GENERATIVE AI AND AI-ASSISTED TECHNOLOGIES IN THE WRITING PROCESS

During the preparation of this work, the authors used GPT-5 in order to edit the manuscript for grammar and clarity. After using this tool, the authors reviewed and edited all content as needed, and take full responsibility for the content of the publication.

---

**Supplementary Figure 1.**
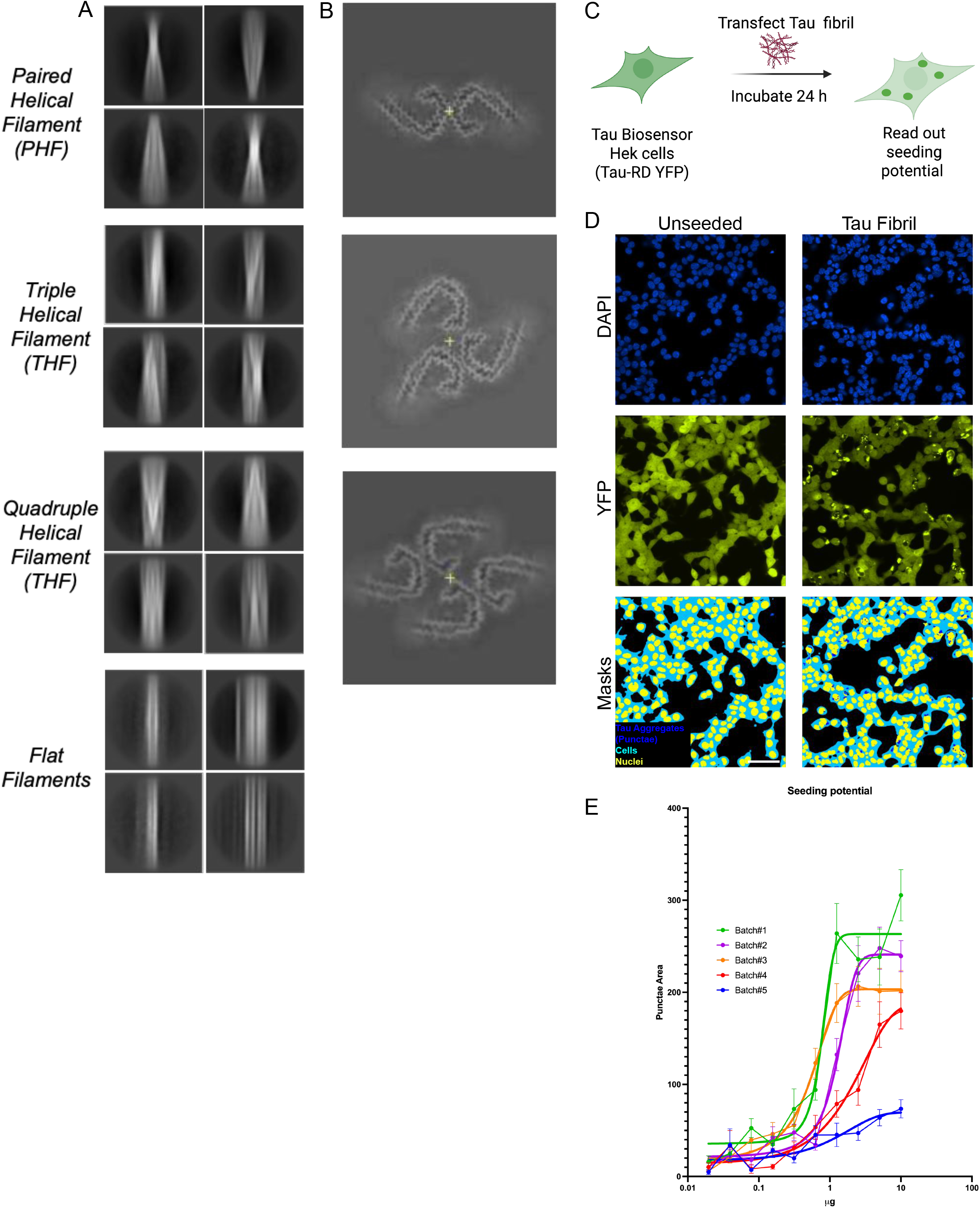
Recombinant Fibrillar Tau (dGAE) Seeding Material. **(A)** Representative CryoEM 2D class averages of recombinant seeding material, including paired, triple, and quadruple helical filaments, and flat filaments. **(B)** Representative central slices of CryoEM 3D reconstructions of recombinant seeding material, including PHFs, THFs, and QHFs. Reconstructions were not accomplished with flat filaments. **(C)** Schematic representation of the HEK biosensor assay. Briefly, HEK biosensor cells were transfected with varying amounts of recombinant Tau fibrils derived from different generated batches and imaged. Seeding potential was expressed through Tau aggregate puncta, which were identified in the YFP channel by masking and quantified accordingly. **(D)** Representative immunofluorescence images of HEK Tau biosensor cells, with and without exposure to Tau fibrils. Scale bar, 100 µm **(E)** Comparison of the seeding potential of the Tau recombinant seeding materials.

**Supplementary Figure 2.**
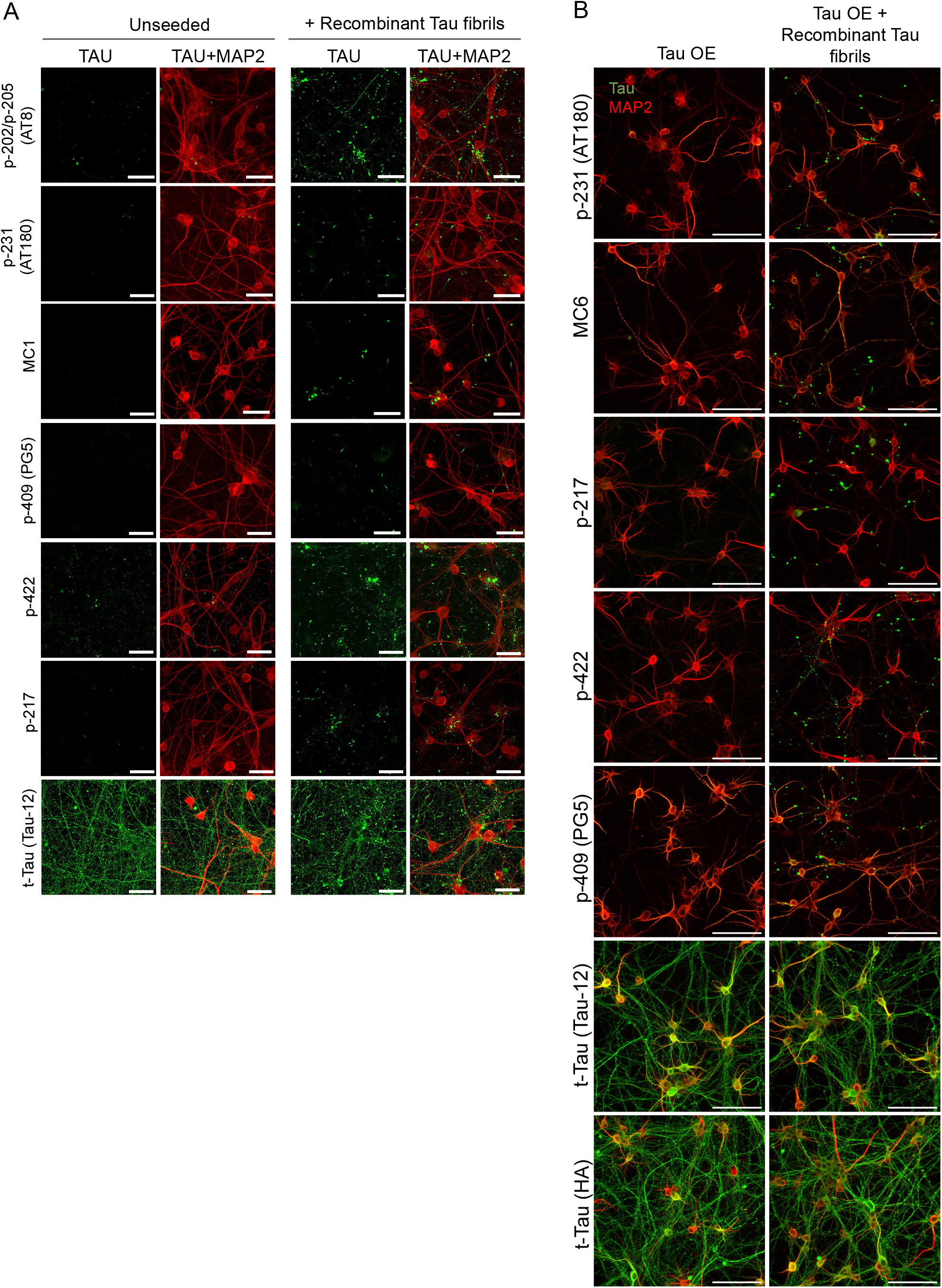
Tau Seeding in NGN2 Neurons Induces Aggregation Labeled with Several p-TAU/misfolded Tau Markers. **(A)** Representative immunofluorescence images of iNGN2 neurons without Tau overexpression seeded with recombinant Tau filaments for 12 weeks, stained with p-202/p-205 (AT8), p-231 (AT180), MC1, p-217, p-422, p-409 (PG5) and t-Tau (Tau-12) antibodies. Scale bars, 50 µm. **(B)** Representative immunofluorescence images of iNGN2 neurons overexpressing Tau and seeded with recombinant Tau filaments for 14 days, stained with p231 (AT180), MC1, p-217, p-422, p-409 (PG5), t-Tau (Tau-12) and t-Tau (HA) antibodies. Scale bars, 100 µm.

**Supplementary Figure 3.**
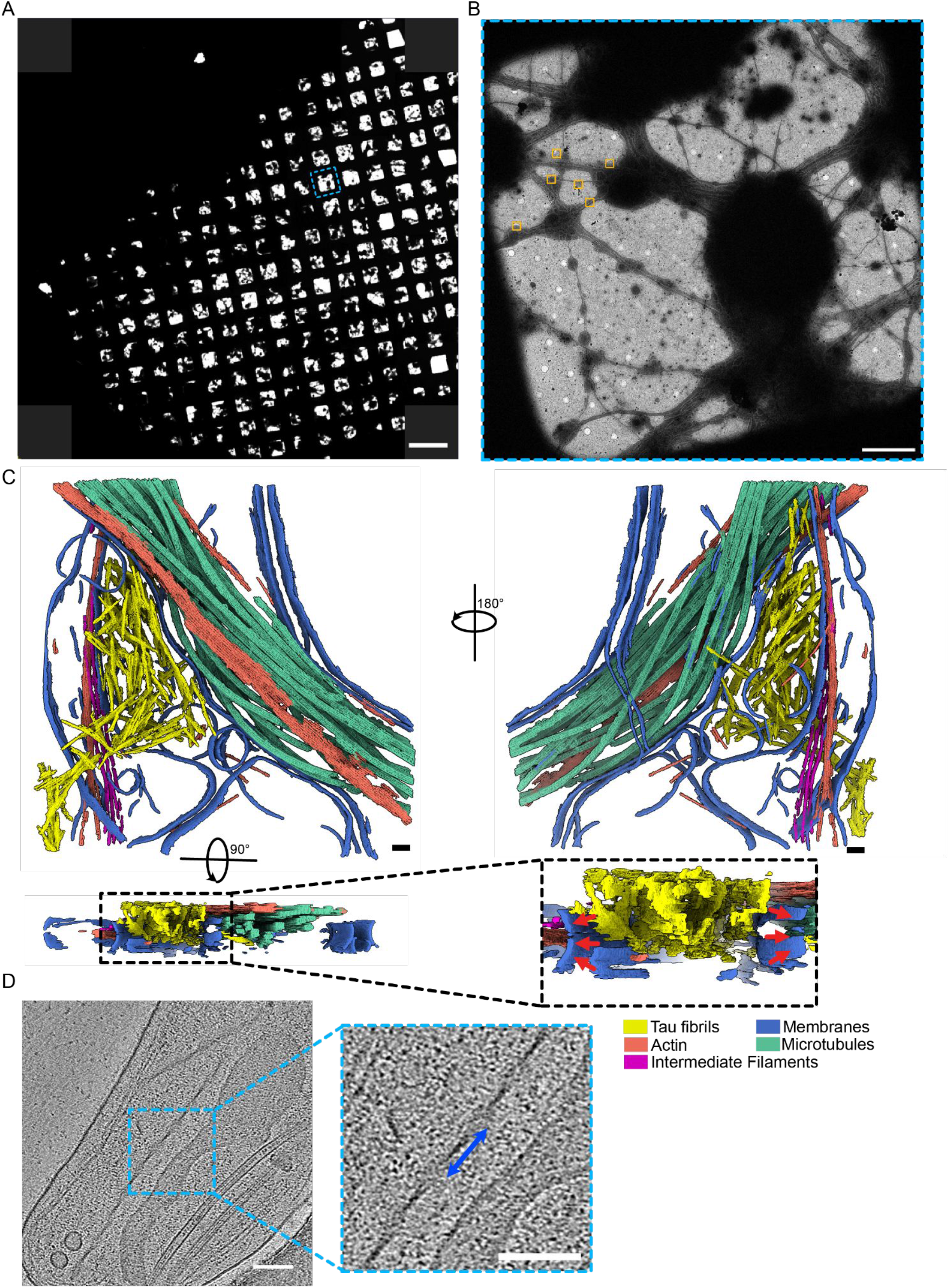
Cryo-ET of iPSC-Derived Neurons Seeded with Tau Fibrils. (A) Atlas of a cryoEM grid, featuring a dense network of iPSC-derived neurons. Scale bar, 0.2 mm. (B) A close-up view of one of the squares outlined in (A) showing an extensive neurite network. Orange boxes indicate example areas of cryo-ET data collection. Scale bar, 10 µm. (C) Orthogonal views of the segmented tomogram showing bundles of short filaments trapped between the membranes and above the cellular structures, in the extracellular space. Red arrows indicate the convex side of the plasma membrane (extracellular space). Scale bars, 100 nm. (D) Example central slices from the tomogram with fibrils whose fold is consistent with the PHF fold. The blue arrow shows the cross-over distance of approximately 80 nm. Scale bars, 100 nm.

**Supplementary Figure 4.**
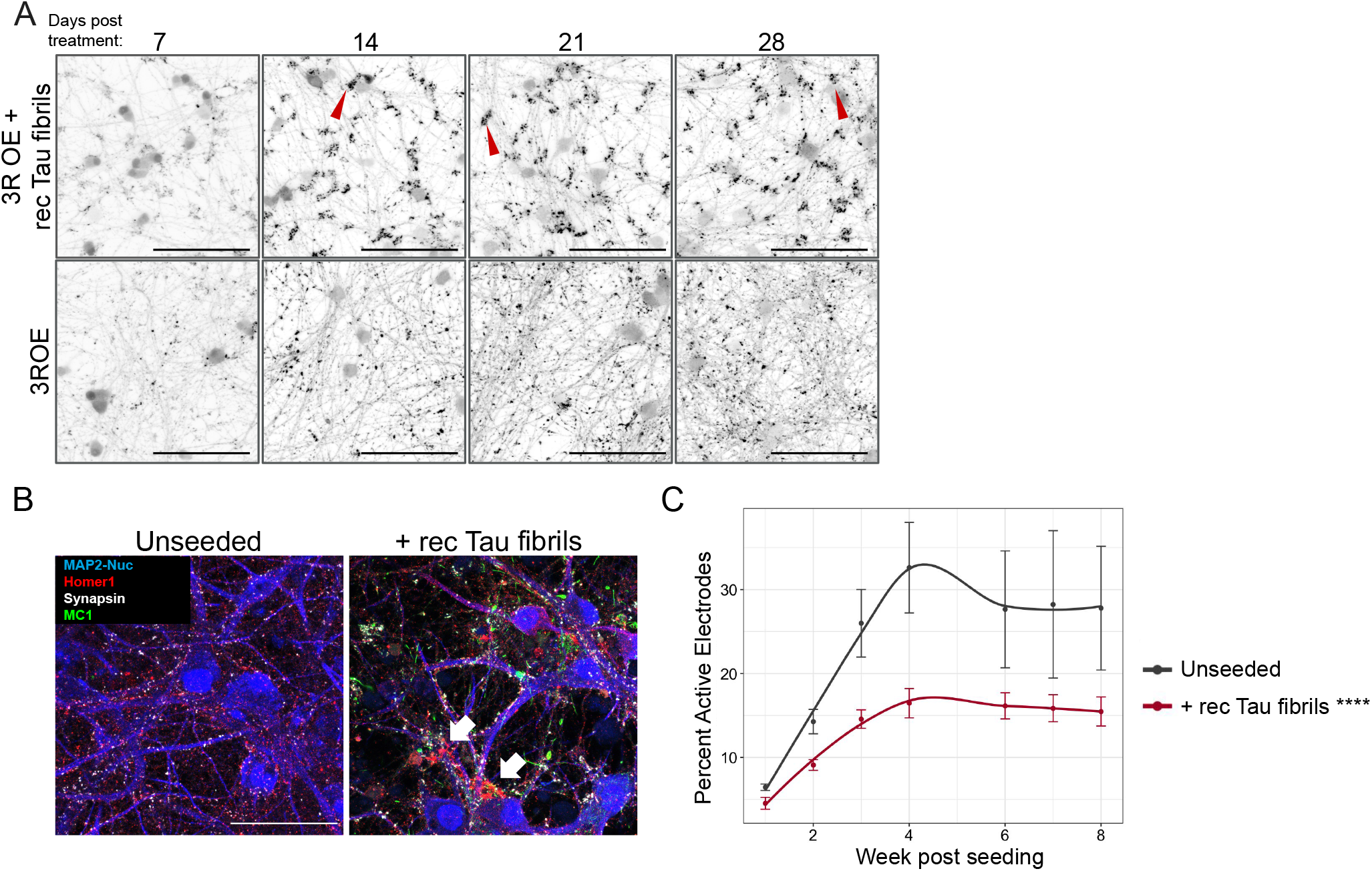
Tau Seeding Causes Progressive Synaptic Vesicle Clustering and Disruption of Network Activity in iNGN2 Neurons. (A) Representative immunofluorescence images of iNGN2 neurons overexpressing Tau, seeded with recombinant Tau filaments, and collected at multiple time points, stained with a synaptophysin antibody. Red arrows mark example synaptic vesicle clusters. Scale bars, 100 µm. **(B)** Representative immunofluorescence images of iNGN2 neurons seeded with recombinant Tau filaments at differentiation day 14 and imaged 12 weeks post seeding. Blue: MAP2-DAPI co-stain; Red: Homer1 (post-synaptic marker); Gray: Synapsin (pre-synaptic marker); Green: MC1 (Tau aggregates). Arrows point to synaptic clumps. Scale bar, 50 µm. (C) High-density multiple electrode array (HD-MEA) readings of neuroactivities, measured by percent active electrodes, in iNGN2 neurons with or without recombinant Tau filaments. Tau seeds were added to neurons at differentiation day 14 and MEA activities were measured weekly post seeding. Recombinant fibril seeding significantly reduces neuronal activities compared with control; n = 3. ****p<0.0001, two-way ANOVA.

**Supplementary Figure 5.**
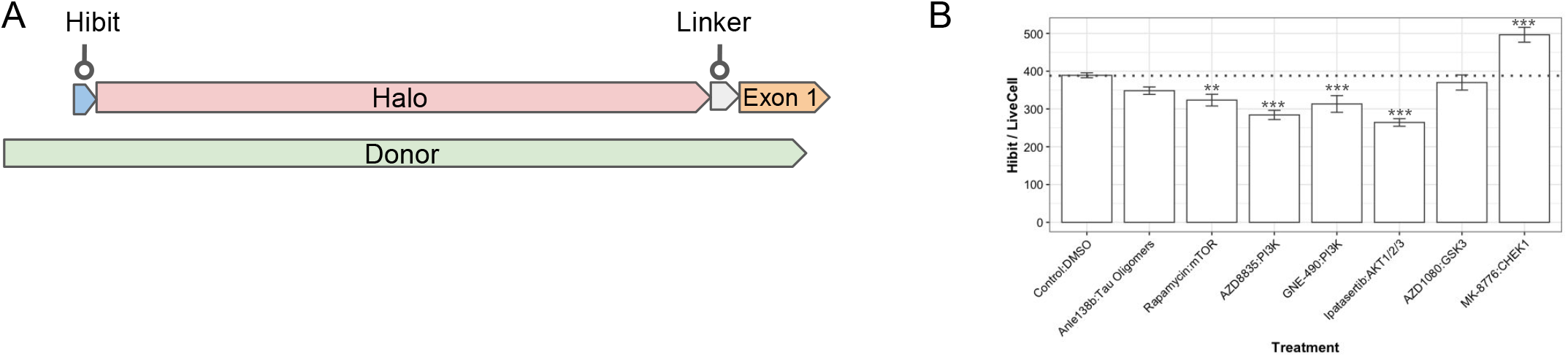
Effects of Selected Compound Hit Reducers on Tau Expression. (A) Schematics on the strategy to knock in a Hibit-Halo tag on the N-terminus of Tau. (B) Normalized level of total Tau measured by Hibit assay intensity over live cell count. Small but significant reductions were found in Rapamycin, AZD8835, GNE490, and Ipatasertip-treated cells. MK8876 significantly increased the total Tau level; n=6 (compounds) or n=21 (DMSO control). *p<0.05, **p < 0.01, ***p<0.001 (one-way ANOVA followed by Dunnett’s test).

